# Specific growth rates and growth stoichiometries of Saccharomycotina yeasts on ethanol as sole carbon and energy substrate

**DOI:** 10.1101/2024.09.27.615414

**Authors:** Marieke Warmerdam, Marcel A. Vieira-Lara, Robert Mans, Jean Marc Daran, Jack T. Pronk

## Abstract

Emerging low-emission production technologies make ethanol an interesting substrate for yeast biotechnology, but information on growth rates and biomass yields of yeasts on ethanol is scarce. Strains of 52 Saccharomycotina yeasts were screened for growth on ethanol. The 21 fastest strains, among which representatives of the Phaffomycetales order were overrepresented, showed specific growth rates in ethanol-grown shake-flask cultures between 0.12 and 0.46 h^-1^. Seven strains were studied in aerobic, ethanol-limited chemostats (dilution rate 0.10 h^-1^). *Saccharomyces cerevisiae* and *Kluyveromyces lactis*, whose genomes do not encode Complex-I-type NADH dehydrogenases, showed biomass yields of 0.59 and 0.56 g_biomass_ g ^-1^, respectively. Different biomass yields were observed among species whose genomes do harbour Complex-I genes: *Phaffomyces thermotolerans* (0.58 g g^-1^), *Pichia ethanolica* (0.59 g g^-1^), *Saturnispora dispora* (0.66 g g^-1^), *Ogataea parapolymorpha* (0.67 g g^-1^), and *Cyberlindnera jadinii* (0.73 g g^-1^). The biomass yield of *C. jadinii*, which also showed the highest biomass protein content of these yeasts, corresponded to 88% of the theoretical maximum achieved when growth is limited by assimilation rather than by energy availability. This study indicates that energy coupling of mitochondrial respiration and its regulation are key factors for selecting and improving yeast strains for ethanol-based processes.

## Introduction

Industrial biotechnology still predominantly uses (partially) refined sugars, such as corn-starch-derived glucose and sugar-cane-derived sucrose, as feedstocks (Nielsen et al., 2013; van Aalst et al., 2022). While sugar-based biotechnological processes can enable lower carbon footprints than petrochemical alternatives (Farrell et al., 2006; Hermann et al., 2007), they are linked with greenhouse-gas (GHG) emissions from agriculture, transport and sugar refining (Salim et al., 2019). In addition, unrestricted expansion of sugar-based industrial biotechnology may interfere with other human needs for arable land (Tilman et al., 2006).

Ethanol, the largest-volume product of industrial biotechnology, is widely used as a ‘drop-in’ transport fuel and increasingly also as renewable chemical building block (Dagle et al., 2020; Fulton et al., 2015). Ethanol production currently relies predominantly on corn-starch hydrolysates or cane sugar (Jacobus et al., 2021). However, advancements in alternative processes increase the potential of ethanol production with reduced GHG emissions.

Biotechnological conversion of agricultural and forestry residues has the potential to substantially reduce GHG emissions of ethanol production (Lynd et al., 2022). While new process modes for such ‘second generation’ ethanol production continue to be explored (Claes et al., 2020), industrial application has already moved to industrial scale (Jansen et al., 2017). Production of ethanol from syngas by acetogenic bacteria (Martin et al., 2016; Phillips et al., 1993) has also been successfully scaled up to industrial level (Heffernan et al., 2020). Although in early phases of development, direct (electro)catalytic production of CO_2_ to ethanol (Birdja et al., 2019) and microbial electrosynthesis (Cabau-Peinado et al., 2024) offer additional perspectives for low-GHG-emission ethanol production.

Total GHG emissions of bioprocesses could be reduced if ethanol obtained from these processes were to be used as feedstock instead of sugars derived from agriculture (van Peteghem et al., 2022a), with the added benefit of industrial biotechnology being less dependent on successful crop harvests. Ethanol is readily consumed by many yeast species (Kurtzman et al., 2011), already a preferred host for industrial biotechnology. Furthermore, concentrated ethanol solutions can be pumped at low temperatures and are self-sterilising, which can contribute to water economy and reduce complexity of large-scale, aerobic processes (Weusthuis et al., 2011). Recent advancements in production of low-GHG-emission ethanol, combined with the potential advantages of ethanol as feedstock for yeast-based industrial biotechnology, have led to interest in finding the ideal host for such bioprocesses. Ethanol has to be respired aerobically for the generation of ATP through the mitochondrial electron transfer system (ETS), indicating that energy conservation in yeast’s metabolism will undoubtedly impact industrial applications.

In yeasts, dissimilation of ethanol starts by its oxidation to acetaldehyde by NAD^+^-dependent alcohol dehydrogenases, after which acetaldehyde is oxidized to acetate by NAD(P)^+^-linked acetaldehyde dehydrogenases. Acetate is then activated to acetyl-coenzyme A (acetyl-CoA) by acetyl-CoA synthetases, a reaction coupled to the hydrolysis of ATP to AMP and pyrophosphate. Because pyrophosphate is subsequently hydrolysed, acetate activation requires a net input of 2 ATP equivalents (Pronk et al., 1994). Complete oxidation of acetyl-CoA to CO_2_ in the tricarboxylic acid cycle yields 3 NADH, 1 quinol (QH_2_) and, through substrate-level phosphorylation, 1 ATP. The overall net ATP yield from ethanol dissimilation depends on the ATP stoichiometry of mitochondrial respiration and is expressed by the P/O ratio. This ratio indicates the number of ATP molecules formed by F_1_F_o_-ATP synthase activity upon transfer of the two electrons from NADH or QH_2_ to oxygen, coupled via proton translocation across the inner mitochondrial membrane by the ETS and subsequent utilisation of the generated proton motive force for the synthesis of ATP molecules. For ethanol dissimilation, the net ATP yield is equal to 5·P/O_NADH_ + 1·P/O_QH2_ – 1 mol ATP (mol ethanol)^-1^, with the caveat that P/O_NADH_ may depend on whether NADH is generated in the yeast cytosol or in the mitochondrial matrix (Verduyn et al., 1991).

The P/O ratio is determined by the composition of the mitochondrial ETS and, when different branches to and from the Q junction can be expressed, by the *in vivo* distribution of electrons over different branches. Model organism *Saccharomyces cerevisiae* has a relatively simple ETS in which electrons from NADH and succinate are both donated to the Q pool (Bakker et al., 2001). *S. cerevisiae* lacks a quinol oxidase (alternative oxidase, AOX) that, in some other yeasts, can bypass the cytochrome c-dependent pathway and the associated proton-translocating sites (Guerrero-Castillo et al., 2012; Joseph-Horne et al., 2001). In *S. cerevisiae*, *in-vivo* P/O_NADH_ and P/O_QH2_ ratios are both approximately 1 (Postmus et al., 2011; Verduyn et al., 1991), resulting in a net ATP yield of approximately 5 mol ATP (mol ethanol)^-1^. It is relevant to note that nuclear and mitochondrial genomes of many non-*Saccharomyces* yeasts harbour the genetic information to encode a multi-subunit mitochondrial proton-pumping NADH dehydrogenase complex (‘Complex I’) (Nosek & Fukuhara, 1994). Involvement of Complex I in mitochondrial NADH oxidation enables higher P/O_NADH_ ratios and thereby, could contribute to higher biomass and product yields on ethanol. However, Complex-I-encoding genes often occur in combination with orthologs of genes encoding the non-proton-pumping *S. cerevisiae* NADH dehydrogenases (Melo et al., 2004), which implies that, depending on regulation of gene expression, electron transfer systems for NADH oxidation in these yeasts may be branched.

Despite its potential as a carbon source for ethanol-based industrial biotechnology, information on specific growth rates and biomass yields of different yeast species on ethanol is limited. The goal of this study is to obtain an indication of the diversity of specific growth rates and biomass yields of Saccharomycotina yeasts when grown on ethanol as sole carbon and energy source. To this end, strains of 52 Saccharomycotina yeasts were screened and those tested positive for growth were assessed semi-quantitatively in microtiter plates for their growth rates on ethanol. Specific growth rates of 21 strains with the fastest growth in microtiter plates were measured in shake-flask cultures. Finally, seven strains, the genomes of five of which harboured the genetic information to encode a Complex-I NADH dehydrogenase, were grown in ethanol-limited chemostat cultures and analysed for their biomass yield, macromolecular composition and energetic efficiency.

## Materials & Methods

### Strains and maintenance

Yeast strains used in this study are listed in Table S1 (Supporting Information). *Saccharomyces cerevisiae* CEN.PK113-7D and *Yarrowia lipolytica* W29 were provided by Dr. P. Kötter (J.-W. Goethe Universität, Frankfurt) and Dr. J.M. Nicaud (Micalis Institute, INRAE, AgroParisTech, Université Paris-Saclay), respectively. *Komagataella phaffii* was obtained as *Pichia pastoris* X-33 from Invitrogen (Thermo Fisher Scientific). All other strains were obtained from the Westerdijk Institute (Utrecht, the Netherlands). For maintenance, strains were grown in shake flasks on YPD medium. YPD was prepared by autoclaving a solution of 10 g L^-1^ yeast extract and 20 g peptone g L^-1^ (Thermo Fisher Scientific) in demineralized water at 121 °C for 20 min. A sterile 50% (w/v) glucose solution, autoclaved separately at 110 °C for 20 min, was then aseptically added to achieve a glucose concentration of 20 g L^-1^. Sterile glycerol was added to stationary-phase cultures at a final concentration of 30% (v/v) and aliquots were stored at -80 °C.

### Synthetic growth media

Synthetic medium (SM) for shake-flask and chemostat cultivation was prepared as described by Verduyn et al. (1992) and contained per litre: 5 g (NH_4_)_2_SO_4_, 3 g KH_2_PO_4_, 0.5 g MgSO_4_·7 H_2_O, 1 mL trace metals solution and 1 mL vitamins solution. The trace elements solution (1000x final medium concentration), containing 4.5 g CaCl_2_·2 H_2_O, 4.5 g ZnSO_4_·7 H_2_O, 3 g FeSO_4_·7 H_2_O, 1 g H_3_BO_3_, 0.84 g MnCl_2_·2 H_2_O, 0.4 g NaMoO_4_·2 H_2_O, 0.3 g CoCl_2_·6 H_2_O, 0.3 g CuSO_4_·5 H_2_O, 0.1 g KI and 15 g EDTA per litre, was added to a solution of the other mineral salts prior to autoclaving at 121 °C for 20 min. For shake-flask cultivation, the pH was adjusted to 6.0 with 2 M KOH prior to autoclaving. For chemostat cultivation, 0.2 mL L^-1^ Pluronic 6100 PE antifoam (BASF, Ludwigshafen, Germany) was added prior to autoclaving. Ethanol and filter-sterilized vitamin solution (1000× final medium concentration, per litre: 50 mg biotin, 1 g Ca-pantothenate, 1 g nicotinic acid, 25 g myo-inositol, 1 g thiamine·HCl, 1 g pyridoxine·HCl, 0.2 g *p*-aminobenzoic acid) were added aseptically after autoclaving. Unless otherwise indicated, ethanol was added to a final concentration of 7.5 g L^-1^ for shake-flask cultivation and 3.75 g L^-1^ for chemostat cultivation. Microtiter plate cultures and inocula for these cultures were grown on buffered synthetic medium described by Jensen et al. (2014) to postpone growth deceleration due to acidification. Compared with SM, this medium contained higher concentrations of (NH_4_)_2_SO_4_ (7.5 g L^-1^), KH_2_PO_4_ (14.4 g L^-1^) and trace elements (2 mL trace metals solution per L). For microtiter plate cultures, 3 g L^-1^ ethanol was added.

### Cultivation techniques

Shake-flask cultures were grown at 30 °C in round-bottom flasks with a working volume of 20% of their nominal capacity and shaken at 200 rpm in an Innova incubator (Eppendorf Nederland BV, Nijmegen, the Netherlands). Late-exponential or early-stationary phase cultures were used to inoculate microtiter plates and chemostats.

Microtiter plate cultures were grown at 30 °C in an image-analysis-based Growth-Profiler system (EnzyScreen BV, Heemstede, the Netherlands) equipped with 96-well plates. Culture volume was 250 µL, plates were agitated at 250 rpm and imaged at 30 min intervals. Wells were inoculated at an OD_660_ of approximately 0.1. Green values of imaged wells were corrected for well-to-well variation by calibration with a 96-well plate containing an *S. cerevisiae* CEN.PK113-7D cell suspension with an externally measured OD_660_ of 2.5. The time to reach stationary phase (TS) was determined as t_2_-t_1_, with t_1_ indicating the time (h) at which the corrected green value was twice as high as the value at inoculation (average of the first three measured green values after inoculation), and t_2_ indicating the first time point at which growth deceleration was observed (Fig. S1, Supporting Information).

Ethanol-limited chemostat cultures were grown at 30 °C and at a dilution rate of 0.10 h^-1^ in 1.5 L bioreactors (Getinge-Applikon, Delft, the Netherlands) with a 1-L working volume. The culture pH was maintained at 5.0 by automatic addition of 2 M KOH, controlled by an ez-Control module (Getinge-Applikon). The working volume was kept at 1.0 L by an electrical level sensor that controlled the effluent pump. The precise working volume of chemostat cultures was measured at the end of each experiment by weighing the broth. Cultures were stirred at 800 rpm and sparged with dried, compressed air (0.5 L min^-1^). Off-gas flow rates from bioreactors were measured with a Mass-Stream Thermal Mass Flow Meter (Bronkhorst, Vlaardingen, the Netherlands). Off-gas was cooled to 4 °C in a condenser to minimise evaporation and dried with a Perma Pure dryer (Inacom Instruments, Veenendaal, the Netherlands). Concentrations of CO_2_ and O_2_ in the inlet and dried exhaust gas were measured with an analyser (Servomex MultiExact 4100, Crowborough, United Kingdom). Cultures were assumed to have reached steady state when, after a minimum of 5 volume changes, CO_2_ production rate, biomass concentration, and high-performance liquid chromatography (HPLC) measurements changed by less than 5% over two consecutive volume changes.

## Analytical methods

Optical density of cultures was measured at 660 nm with a Jenway 7200 spectrophotometer (Jenway, Staffordshire, United Kingdom) after dilution with demineralised water to OD_660_ values between 0.1 and 0.3. Specific growth rates of shake-flask cultures were calculated by linear regression of the relation between ln(OD_660_) and time in the exponential growth phase, including at least 6 time points over at least 3 biomass doublings.

For biomass dry weight determination, duplicate culture samples of exactly 10.0 mL (or 5.00 mL for *Ogataea parapolymorpha*) were filtered over pre-dried and pre-weighed membrane filters (0.45 μm pore size; Pall Corporation, Ann Arbor, MI). Filters were then washed with demineralized water, dried in a microwave oven at 320 W for 20 min, and immediately reweighed (Postma et al., 1989).

Metabolite concentrations in culture supernatants and media were measured with an Agilent 1260 Infinity HPLC system (Agilent Technologies, Santa-Clara, CA) equipped with a Bio-rad Aminex HPX-87H ion exchange column (Bio-Rad, Hercules, CA), operated at 60 °C with 5 mM H_2_SO_4_ as mobile phase at a flow rate of 0.600 mL min^-1^. Culture supernatant was obtained by centrifuging 1 mL aliquots at 13,000 rpm for 5 min with a Biofuge Pico centrifuge (Heraeus Instruments, Hanau, Germany). Metabolite concentrations in steady-state chemostat cultures were analysed after rapid quenching of culture samples with cold steel beads (Mashego et al., 2003).

Whole-cell protein content was determined essentially as described by Verduyn et al., (1990). Exactly 20.0 mL of culture (2 - 4 g L^-1^ dry weight) was harvested from bioreactors by centrifugation (5 min, 10,000 g at 4°C) in an Avanti J-E centrifuge (Beckman Coulter Life Sciences, Indianapolis, IN), using a JA-25.50 rotor. The pellet was washed once, and the final pellet was resuspended in water to a final volume of 10.0 mL and stored at -20 °C. After thawing at room temperature, samples were analysed with the alkaline copper sulphate method, using bovine serum albumin (Sigma-Aldrich, St. Louis, MO) as a standard.

Analysis of whole-cell lipid content was based on the method described by Izard & Limberger, (2003). Phosphoric acid–vanillin reagent was prepared by adding 0.120 g of vanillin (Sigma-Aldrich) to 20 mL of demineralised water and adjusting the volume to 100 mL with 85% H_3_PO_4_. Biomass samples from bioreactors were obtained as described for whole-cell protein content determination. After thawing at room temperature, 100 µL of the cell suspension was added to a stoppered glass tube, together with 1 mL of concentrated H_2_SO-_4_. Samples were boiled for 10 min and subsequently cooled for 5 min at room temperature in a water bath. Then, 2.5 mL of phosphoric acid-vanillin reagent were added, and samples were incubated for 15 min at 37 °C. Samples were cooled for 10 min in a water bath at room temperature. Absorbance at 530 nm was measured with a Jenway 7200 spectrophotometer against a demineralised water sample. Lipid contents were calculated based on a dilution range of sunflower oil (Sigma-Aldrich) in chloroform. Prior to H_2_SO_4_ addition, glass tubes containing the standard solutions were placed in a water bath at 70 °C until chloroform was fully evaporated, after which 100 µL of demineralised water was added for volume correction.

Whole-cell RNA content was quantified by determining absorbance at 260 nm of RNA degraded by alkali and extracted from the culture in HClO_4_ (adapted from Beck et al., (2018) and Benthin et al., (1991)). Samples of 2.00 mL (4 - 8 mg biomass dry weight) were harvested from bioreactors and centrifuged at 4,000 rpm for 10 min at 4 °C in an Eppendorf 5810 R (Eppendorf Netherlands BV, Nijmegen, the Netherlands). Pellets were washed once with demineralised water and, after the second round of centrifugation, stored at -20 °C. Pellets were thawed at room temperature and washed three times with 3 mL cold 0.7 M HClO_4_ and digested with 3 mL 0.3 M KOH for 60 min in a water bath set at 37 °C, while vortexing briefly every 15 min. After addition 1 mL of 3 M HClO_4_, supernatant was collected and precipitates were washed twice with 4 mL cold 0.5 M HClO_4_. The collected combined supernatant was centrifuged to remove KClO_4_ precipitates. Absorbance at 260 nm was measured in quartz cuvettes using a Hitachi U-3010 spectrophotometer (Sysmex Europe GmbH, Norderstedt, Germany) against a standard prepared with RNA from yeast (Sigma-Aldrich).

Whole-cell total carbohydrate contents were quantified with a phenol-sulfuric acid-based method (based on Herbert et al. (1971) and Lange & Heijnen (2001)). Briefly, 1.00 mL (2 – 4 mg biomass dry weight) samples from bioreactor cultures were harvested and frozen as described for whole-cell RNA analysis. Frozen samples were thawed on ice and resuspended in 10 mL demineralised water. 5 mL of concentrated H_2_SO_4_ was added to a mixture of 1 mL of resuspended sample and 1 mL of aqueous phenol solution (50 g L^-1^) in a stoppered glass tube. After boiling for 15 min and subsequent cooling to room temperature, absorbance at 488 nm was measured. Samples were measured against a 1:2 mannose:glucose standard. Results were corrected for presence of nucleic pentoses by using extinction coefficients of 0.36 and 0.26 for g L^-1^ RNA and DNA, respectively, and using the RNA content determined as described above and assuming a constant DNA cellular content of 0.5 wt-% (Lange & Heijnen, 2001).

### Phylogenetic tree construction

A phylogenetic tree of the studied yeast species was constructed by the Neighbour-Joining method (Saitou & Nei, 1987). A bootstrap consensus tree inferred from 500 replicates was taken to represent the genealogy of the analysed taxa (Felsenstein, 1985). Evolutionary distances were computed with the p-distance method (Nei & Kumar, 2000) applied to LSU D1/D2 nucleotide sequences deposited in the Yeast IP database (Weiss et al., 2013). When the LSU D1/D2 sequence of a strain used in this study was not available, sequences of the type strain for the species were used instead. Ambiguous positions were removed for each sequence pair, resulting in a dataset with 635 positions in total. Evolutionary analyses were conducted in MEGA11 (Tamura et al., 2021).

### Genomic analysis for genes involved in the synthesis of respiratory chain components

To infer presence and absence of genes encoding specific components of the mitochondrial respiratory chain, protein sequences of *Neurospora crassa* deposited at the National Center for Biotechnology Information (NCBI, https://www.ncbi.nlm.nih.gov/) or of the *S. cerevisiae* CEN.PK strain deposited at the Saccharomyces genome database (SGD, https://www.yeastgenome.org/) were used as queries in a tBLASTn search on the NCBI website using the default settings (Table S2, Supporting Information). The reference sequence as denoted by NCBI was used when multiple genomic datasets were available.

### Stoichiometric analysis with a core metabolic model

Growth stoichiometries of aerobic, ethanol-grown yeast cultures were analysed with a core metabolic network model based on *S. cerevisiae* as described by Daran-Lapujade et al., (2004). The stoichiometry matrix representing the core metabolic network (df=2) consisted of 161 metabolites and 154 reactions divided over three compartments (Supporting Information 1). The matrix was solved assuming a steady state (S·v=0) with the *rref* function in MATLAB R2021b, expressing all metabolic fluxes as a function of growth rate and maintenance.

## Data analysis

Time to stationary phase, specific growth rates and corresponding standard deviations were calculated using Microsoft Excel. All other statistical analyses were performed with GraphPad Prism version 10.1.2.

## Results

### Semi-quantitative comparison of growth kinetics on ethanol of a selection of budding yeast strains

To assess diversity of ascomycete yeasts with respect to their specific growth rates on synthetic medium with ethanol, strains of 52 species covering multiple phylogenetic clades (Shen et al., 2018, Fig. 1) were selected for characterization in an automated, image-analysis-based growth profiler set-up (Duetz, 2007). The selected species had previously been scored positive for growth on ethanol in taxonomic studies (Kurtzman et al., 2011). However, 8 of the 52 selected strains (Table S1, Supporting Information) did not show significant growth on ethanol after >3 days incubation and were eliminated from further characterisation (*Dekkera bruxellensis* CBS 2499, *Kazachstania africana* CBS 2517, *Naumovozyma castellii* CBS 4309, *Priceomyces haplophilus* CBS 2028, *Saccharomyces arboricola* CBS 10644, *Maudiozyma bulderi* CBS 8638, *Saccharomyces kudriavzevii* CBS 8840, and *Zygotorulaspora mrakii* CBS 4218). Three additional strains were excluded because they flocculated in ethanol-grown shake-flask pre-cultures (*Eremotecium cymbalariae* CBS270.75, *Lachancea kluyveri* CBS 3082, and *Lachancea thermotolerans* CBS 6340). Growth-profiler experiments with the remaining 41 strains were performed at 30 °C with synthetic medium supplemented with 3 g L^-1^ ethanol. Since image-analysis by the Growth Profiler set-up yields a non-linear, strain-dependent relation between biomass concentration and read-out, time-to-stationary-phase (TS) was used as a proxy for growth rate (Fig. S1, Supporting Information). The 41 strains showed TS values ranging from 5.7 ± 0.3 h for *Cyberlindnera jadinii* CBS 621 to 65 ± 4 h for *Saccharomyces mikatae* CBS 8839 (Fig. 1 and S2, Supporting Information). Despite showing growth on ethanol in pre-cultures, *Zygosaccharomyces rouxii* CBS 732, *Vanderwaltozyma polyspora* CBS 2163, *Tetrapisispora phaffii* CBS 4417, and *Ogataea methylivora* CBS 7300 showed no or extremely slow growth on ethanol in growth-profiler experiments (Fig. S2 and Table S1, Supporting Information). Fast growth (TS values of 8 ± 2 h) was observed in all tested yeasts strains that belonged to the Phaffomycetales order (n=8, genera *Barnettozyma, Cyberlindnera*, *Phaffomyces*, *Starmera*, and *Wickerhamomyces*).

**Figure 1.**
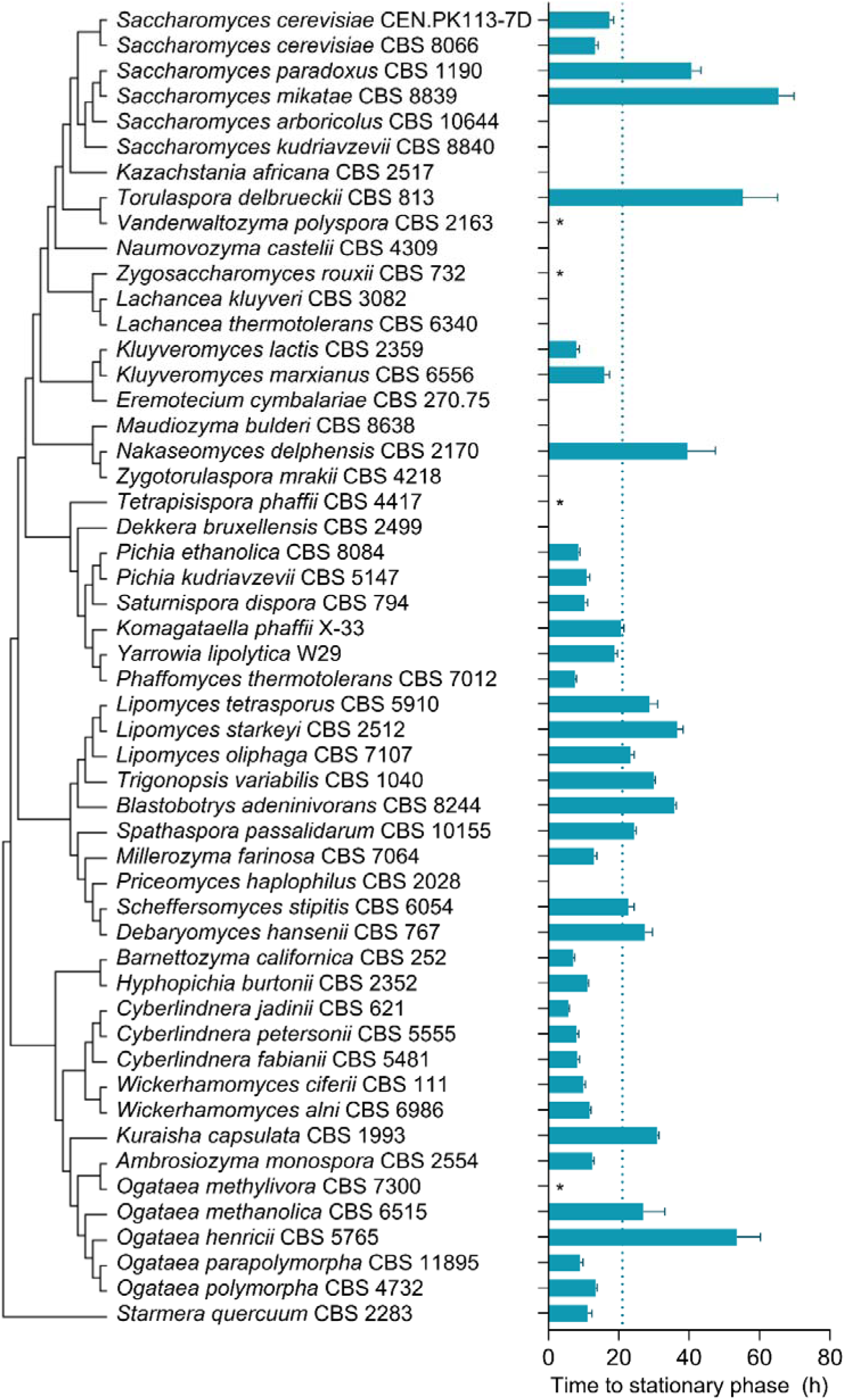
Time required to reach stationary phase (TS) on synthetic medium with 3 g L^-1^ ethanol as sole carbon source of strains of different Saccharomycotina species. The data were obtained from batch cultures in microtiter plates grown at 30 °C and at an initial pH of 6.0. Time to stationary phase was determined as the interval between the time point at which the green value measured by the Growth Profiler setup had doubled relative to its value immediately after inoculation, and the first time point at which a deceleration of the increase of the green value was observed (Fig. S1 and S2, Supporting Information). Bars represent average and standard deviation of at least four independent experiments for each yeast strain. For strains denoted with an asterisk (*) a doubling of the green value was not observed within the timeframe of the experiment (196 h). For strains without annotation, growth characteristics observed in inoculum shake flasks were the reason for exclusion from MTP screening (Table S1, Supporting Information). Strains with a TS < 21 h (cut-off value indicated with dotted line) were further characterised in shake-flask batch cultures (Fig. 2). Phylogenetic relationships are represented as a bootstrap consensus tree (500 replicates) inferred with the Neighbour-Joining method based on LSU D1/D2 sequences deposited in the Yeast IP database (Weiss et al., 2013).

### Comparison of specific growth rates on ethanol in shake-flask cultures

The 21 yeast strains that showed a TS below 21 h in the growth-profiler experiments were grown in shake-flask cultures for an accurate measurement of their specific growth rates on ethanol (Fig. 2). Specific growth rates of these strains on SM with 7.5 g L^-1^ ethanol ranged from 0.12 ± 0.00 h^-1^ for *Komagataella phaffii* X-33 to 0.46 ± 0.01 h^-1^ for *Pichia kudriavzevii* CBS 5147 (Fig. 2 and S3, Table S1, Supporting Information), corresponding to doubling times of 5.8 h and 1.5 h, respectively. As anticipated, specific growth rates measured in shake-flask cultures negatively correlated with growth-profiler TS values (Fig. S4, Supporting Information). Outliers coincided with atypical growth characteristics in shake-flask cultures. For example, after inoculation, shake-flask cultures of *P. kudriavzevii* CBS 5147 showed an initial fast decrease in their optical density which was followed by a short, extremely fast exponential growth phase (Fig. S3, Supporting Information). Other ‘outlier’ strains showed a prolonged lag phase in shake-flask cultures and one exhibited pseudohyphal growth (i.e. *Ambrosiozyma monospora* CBS 2554), which precluded accurate growth-rate measurements (Table S1, Supporting Information).

**Figure 2.**
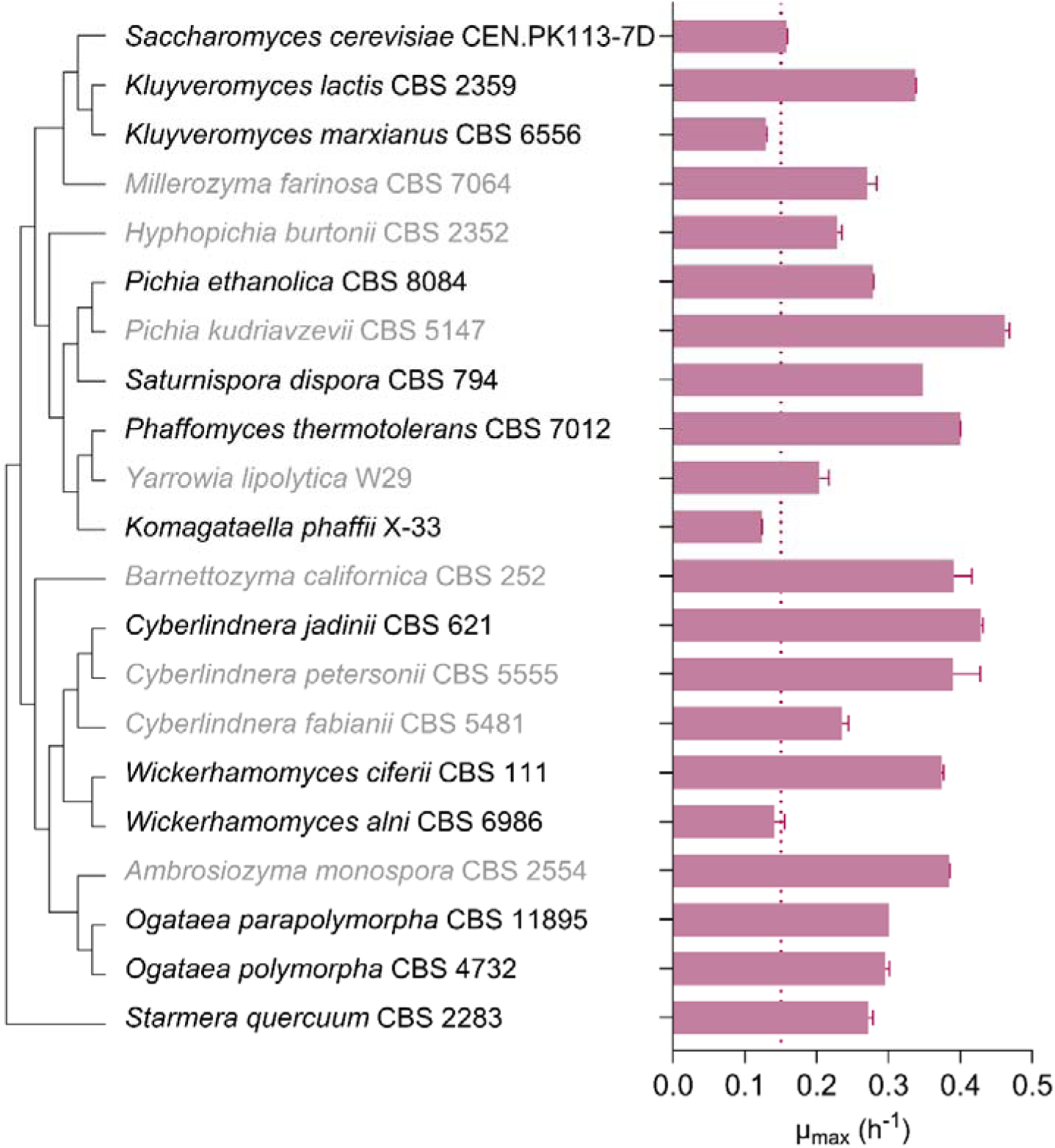
Specific growth rates (µ_max_) of strains of selected Saccharomycotina species, calculated from optical density (OD_660_) measurements during the exponential growth phase of shake-flask batch cultures on synthetic medium with 7.5 g L^-1^ ethanol as sole carbon source. Cultures were grown at 30 °C and at an initial pH of 6.0. Bars show averages with standard deviations of specific growth rates calculated from independent duplicate experiments. Strains with µ_max_ < 0.15 h^-1^ (dotted line) were excluded from further characterisation. Strains with names displayed in grey showed irregular growth characteristics and were also excluded from further characterization (Fig. S3 and Table S1, Supporting Information). Phylogenetic relationships are represented as a bootstrap consensus tree (500 replicates) inferred with the Neighbour-Joining method based on LSU D1/D2 sequences deposited in the Yeast IP database (Weiss et al., 2013).

Based on the results from previous stage of the screening, i.e. the Growth-profiler experiments (Figure 1), all tested strains belonging to the Phaffomycetales order were selected for shake-flask characterisation. Within the subgroup of faster-growing species that were selected for shake-flask characterisation, the performance in terms of high growth rate of the Phaffomycetales strains was less pronounced. The average µ_max_ of the Phaffomycetales strains was 0.33 ± 0.11 h^-1^, whilst the average µ_max_ of all 21 strains tested in shake flasks was 0.29 ± 0.10 h^-1^. This difference is not statistically significant (unpaired two-tailed t test, p = 0.42), unlike comparing the average TS of the Phaffomycetales strains (8 ± 2 h) to the average of the 37 strains where TS could be quantified in the previous stage of the screening (21 ± 15 h, p = 0.02).

### Selection of strains with different respiratory-chain composition based on genome data

Composition of the mitochondrial electron transfer system can significantly affect growth energetics of respiring yeast cultures. NADH oxidation by a mitochondrial proton-pumping NADH dehydrogenase complex (‘Complex I’) enables higher ATP yields from oxidative phosphorylation than NADH oxidation by non-proton-pumping, single subunit mitochondrial NADH dehydrogenases such as Ndi1 in *S. cerevisiae* (Juergens et al., 2020). Conversely, activity of a non-proton pumping alternative oxidase (AOX) can decrease the ATP stoichiometry of mitochondrial respiration (Guerrero-Castillo et al., 2012; Joseph-Horne et al., 2001). We therefore assessed presence and absence of relevant genes in the genomes of 10 yeast species that showed specific growth rates on ethanol of at least 0.15 h^-1^ in shake-flask cultures (Fig. 2 and 3, Table S2, Supporting Information). Based on this analysis, three subgroups were identified: (i) absence of genes encoding Complex I and AOX (*Saccharomyces cerevisiae*, *Kluyveromyces lactis*, and *Starmera quercuum*), (ii) presence of genes encoding Complex I proteins and AOX (*Pichia ethanolica*, *Saturnispora dispora*, *Phaffomyces thermotolerans*, *Cyberlindnera jadinii*, and *Wickerhamomyces ciferii*), and (iii) presence of Complex I-encoding genes but absence of AOX genes (*Ogataea parapolymorpha* and *Ogataea polymorpha*). All three subgroups contained genetic homology with genes known to encode cytochrome c, subunits of ubiquinone:cytochrome c oxidoreductase (Complex III) and cytochrome c oxidase (Complex IV), as well as non-proton-translocating, single-subunit NADH dehydrogenases. *S. cerevisiae* CEN.PK113-7D and *K. lactis* CBS 2359 (both subgroup i), *P. ethanolica* CBS 8084, *S. dispora* CBS 794, *P. thermotolerans* CBS 7012, and *C. jadinii* CBS 621 (subgroup ii, the latter two belonging to the Phaffomycetaceae family), and *O. parapolymorpha* CBS 11895 (subgroup iii) were selected as representatives for a quantitative analysis of biomass yields and biomass composition in chemostat cultures.

**Figure 3.**
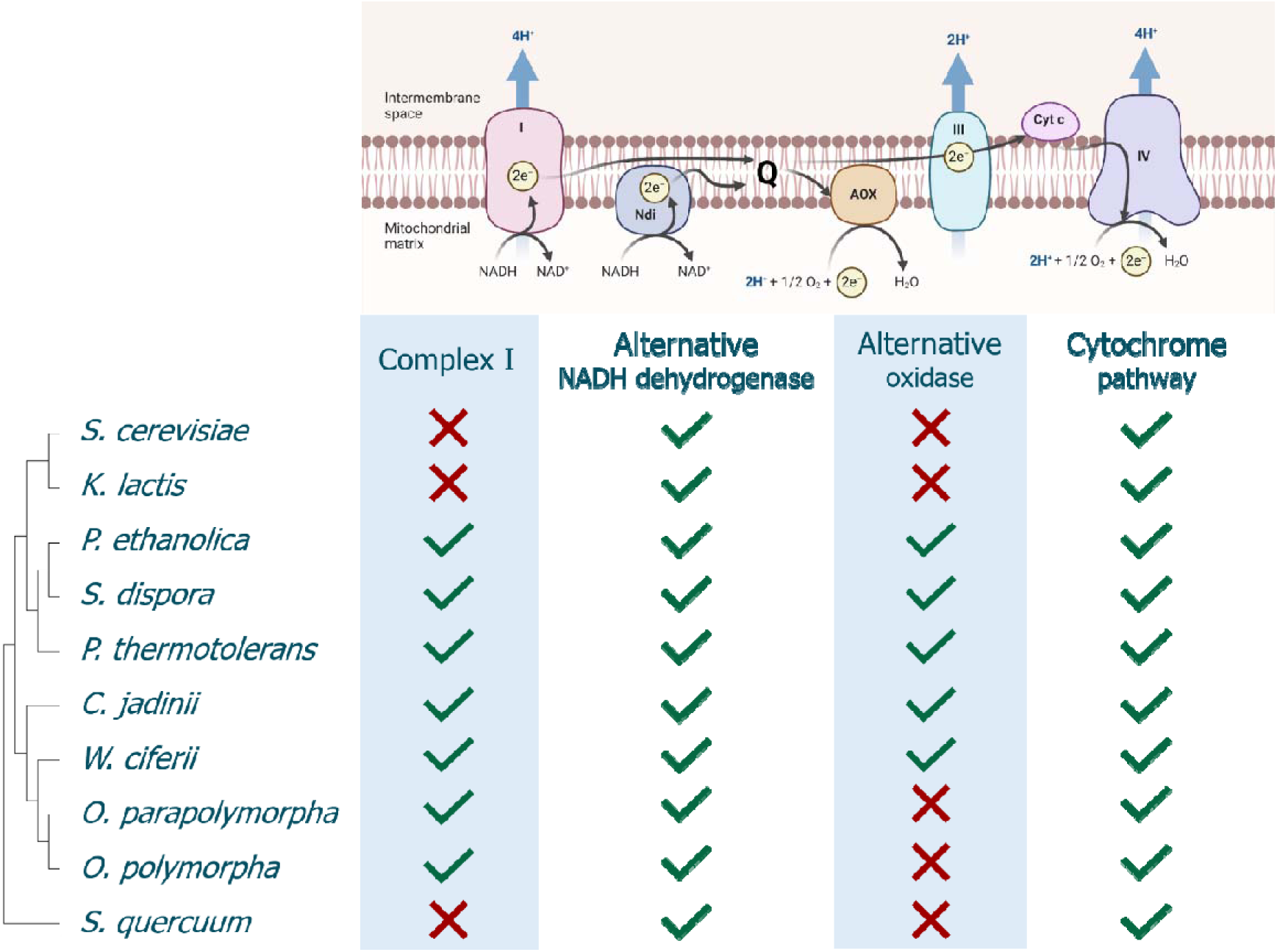
Presence and absence of genes encoding key respiratory chain components in a selection of Saccharomycotina species. Phylogenetic relationships are represented as a bootstrap consensus tree (500 replicates) inferred with the Neighbour-Joining method based on LSU D1/D2 sequences deposited in the Yeast IP database (Weiss et al., 2013). Presence or absence of genes encoding respiratory-chain components was inferred from a tBLASTn search of genomic datasets available in the database of the National Center for Biotechnology Information (NCBI, Table S2, Supporting Information). Abbreviations: (Complex) I NADH:ubiquinone oxidoreductase type 1; Ndi NADH:ubiquinone oxidoreductase type 2; Q quinone; (Complex) III ubiquinone:cytochrome c oxidoreductase; Cyt c cytochrome c; AOX alternative oxidase; (Complex) IV cytochrome c oxidase. Figure created with BioRender.com.

### Biomass yields and biomass composition in ethanol-limited chemostat cultures

In ethanol-limited chemostat cultures, operated at a dilution rate of 0.10 h^-1^ (Fig. 4 and Table 1), biomass yields of the seven selected yeast strains ranged from 0.56-0.73 g_biomass_ g ^-1^ ethanol S. cerevisiae and *K. lactis*, both reported to lack a Complex I NADH dehydrogenase (Nosek & Fukuhara, 1994, Fig. 3), showed similar biomass yields (0.59 ± 0.00 and 0.56 ± 0.00 g g^-1^, respectively). Under the same conditions, higher biomass yields on ethanol were observed for the Complex-I-containing species *S. dispora* (0.66 ± 0.00 g g^-1^), *O. parapolymorpha* (0.67 ± 0.01 g g^-1^) and *C. jadinii* (0.73 ± 0.00 g g^-1^). Although genes encoding Complex-I subunits were also encountered in the genomes of *P. thermotolerans* and *P. ethanolica*, the biomass yield on ethanol of these strains at D = 0.10 h^-1^ (0.58 ± 0.01 g g^-1^ and 0.59 ± 0.00 g g^-1^, respectively) was more similar to that of *S. cerevisiae* and *K. lactis* than to that of *S. dispora*, *O. parapolymorpha* and *C. jadinii* (Fig. S5, Supporting Information). Biomass yields of the seven strains on oxygen ranged from 14.5 ± 0.1 to 22.1 ± 0.4 g dry weight [mol O_2_]^-1^ and, as anticipated, correlated positively with their biomass yields on ethanol (Table 1).

**Figure 4.**
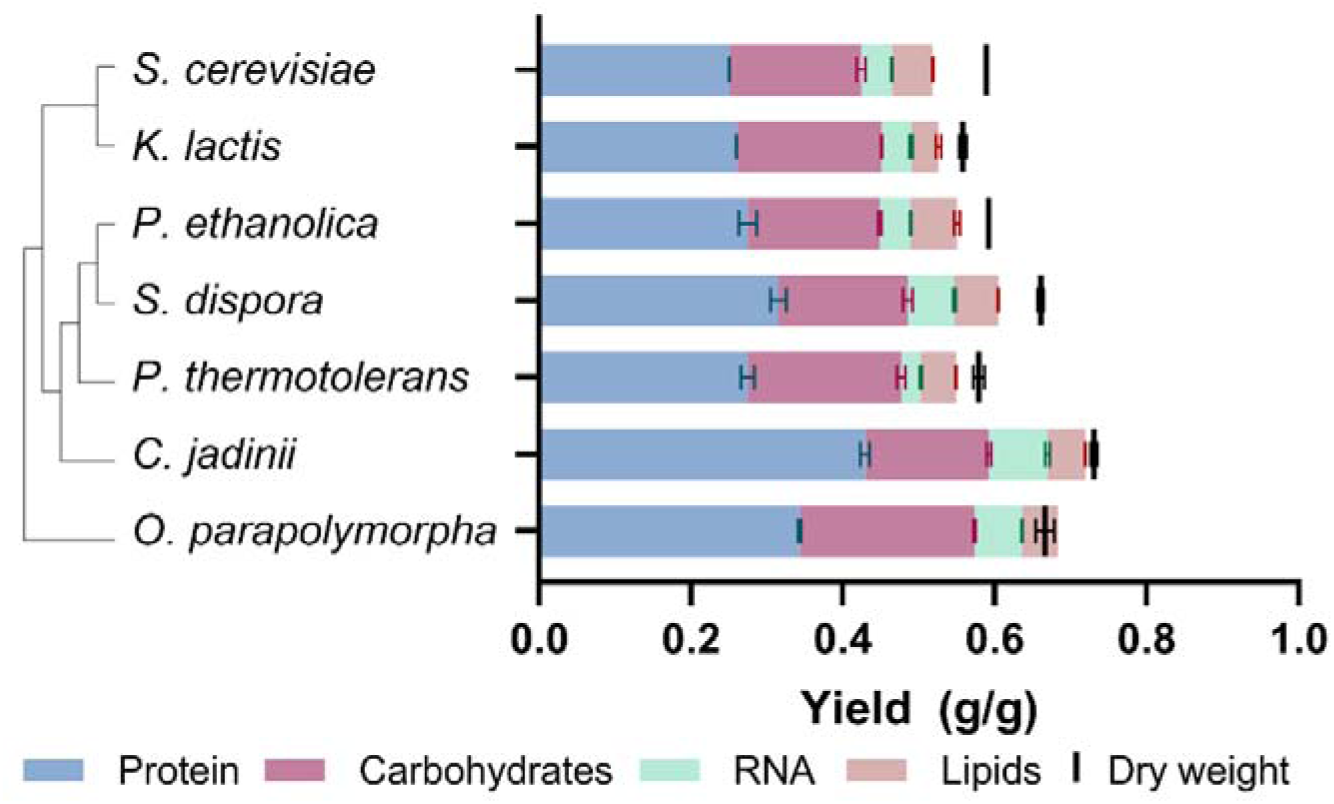
Biomass yield and macromolecular biomass composition of seven different Saccharomycotina species grown in aerobic ethanol-limited chemostat cultures at 30 °C, pH 5.0 and at a dilution rate of 0.10 h^-1^. Data are represented as average and standard deviation of data from independent duplicate chemostat cultures for each yeast strain (Table 1). Phylogenetic relationships are represented as a bootstrap consensus tree (500 replicates) inferred with the Neighbour-Joining method based on LSU D1/D2 sequences deposited in the Yeast IP database (Weiss et al., 2013).

**Table 1.**
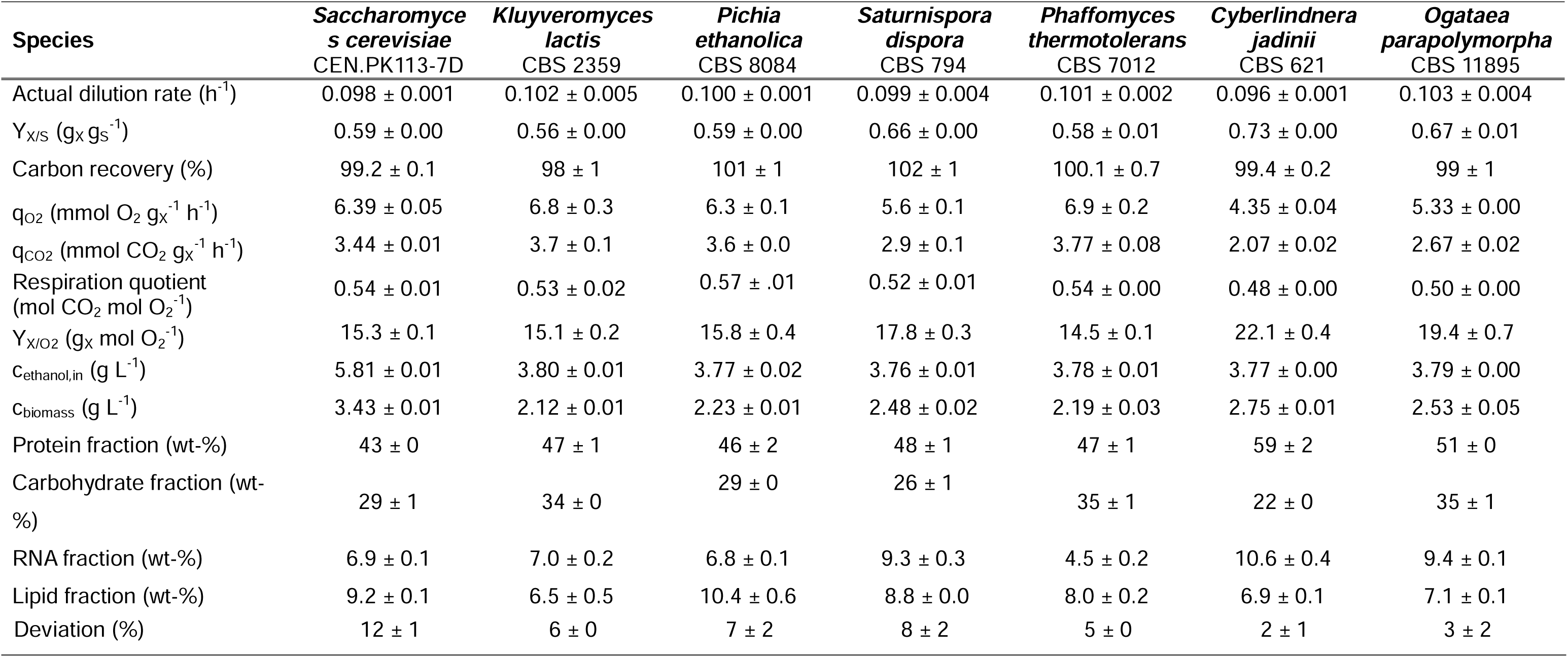
Physiological parameters of yeast species grown in aerobic ethanol-limited chemostat cultures at 30 °C, pH 5.0 and at a dilution rate h^-1^. Ethanol concentrations in the effluent were below the detection limit (3 mg L^-1^). The deviation in the last row indicates the difference een the overall dry weight biomass concentration and the sum of the four biomass components. Averages and standard errors were ned from duplicate experiments.

Because energy costs of protein, carbohydrates, nucleic acids and lipid synthesis are different, comparisons of growth energetics should take into account macromolecular composition of biomass. Therefore, contents of protein, carbohydrates, RNA and lipid were determined for biomass samples taken from chemostat cultures of the seven strains (Fig. 4). Of the four major macromolecular biomass constituents, protein is the most expensive in terms of ATP requirement (Verduyn et al., 1991). Protein content of the biomass of the seven yeasts varied from 43 to 59% (Table 1), with the highest protein content found in *C. jadinii*, which also showed the highest overall biomass yield. Conversely, *C. jadinii* biomass had the lowest carbohydrate content at 22.3 ± 0.3 wt-%, whilst the carbohydrate content for the other species ranged from 26-35 wt-%. The RNA content also varied among the species, with the highest content measured in *C. jadinii* biomass (10.6 ± 0.4 wt-%), and the lowest in *P. thermotolerans* (4.5 ± 0.2 wt-%). The lipid content seemed to have the smallest deviation between the species, ranging from 6.5-9.2 wt-%.

### Theoretical analysis of growth energetics in ethanol-limited chemostat cultures

When expressed per mole of carbon, ethanol (C_2_H_6_O) has a degree of reduction (γ) of 6, while, when ammonium is used as nitrogen source (used as reference γ = 0), the generic elemental biomass composition CH_1.8_O_0.5_N_0.2_ corresponds to γ = 4.2 (Roels, 1980). Assimilation of ethanol into biomass therefore leads to the generation of reducing equivalents (NAD(P)H and QH_2_) whose oxidation by mitochondrial respiration can generate ATP. A theoretical maximum biomass yield is reached when the ATP yield from respiration of the reduced cofactors generated in assimilation matches the ATP requirement in assimilation (Verduyn et al., 1991). In such a situation, assimilation generates enough redox cofactors to meet biosynthetic requirements for ATP and, consequently, complete dissimilation of additional ethanol to CO_2_ and water is not needed.

The theoretical maximum yields of the *S. cerevisiae*, *K. lactis*, *P. ethanolica*, *S. dispora*, *P. thermotolerans*, *C. jadinii*, and *O. parapolymorpha* strains used in the chemostat experiments were calculated with a core stochiometric model of yeast metabolism (Table 2). Molecular and elemental biomass composition were based on the experimentally determined whole-cell contents of protein, RNA, carbohydrate, and lipid of the different yeast species when grown in ethanol-limited steady state chemostats at D = 0.10 h^-1^. DNA content was assumed to be 0.5 wt-% for all strains (Lange & Heijnen, 2001). The oxidation of ethanol and acetaldehyde was modelled to generate cytosolic NADH. For all yeast species, a mechanistic H^+^/O ratio of 6 was assumed for NADH generated in the cytosol and for QH_2_ generated in the mitochondria. For NADH generated in the mitochondrial matrix, H^+^/O ratios of 6 and 10 were assumed for yeasts lacking and containing a Complex I NADH dehydrogenase, respectively. The theoretical maximum biomass scenario was simulated via minimising the H^+^/ATP ratio of the mitochondrial ATP synthase in the model, whilst preventing reverse electron flow predicted by the model as a lower boundary (Table S3, Supporting Information). Growth-rate independent ATP turnover for cellular maintenance and growth-coupled ATP costs were excluded from these theoretical maximum biomass yield simulations.

**Table 2.** Maximum theoretical biomass yields on ethanol as sole substrate, estimated with a stoichiometric model of the core metabolic network of yeast (Supporting Information 1). Elemental and macromolecular biomass compositions were based on measurements of protein, RNA, carbohydrates and lipids in biomass samples from ethanol-limited chemostats (Table 1). DNA content was assumed constant at 0.5-wt% (Lange & Heijnen, 2001). The third column displays for each yeast species the percentage that the experimentally determined yield makes up of the maximum theoretical biomass yield.

| Organism | Biomass yield (g g<sup>-1</sup>) |  | Experimental/<br>Theoretical |
| --- | --- | --- | --- |
|  | Experimental | Theoretical<br>maximum |  |
| *Saccharomyces cerevisiae*<br>CEN.PK113-7D | 0.59 ± 0.00 | 0.826 | 71 % |
| *Kluyveromyces lactis*<br>CBS 2359 | 0.56 ± 0.00 | 0.831 | 67 % |
| *Pichia ethanolica*<br>CBS 8084 | 0.59 ± 0.00 | 0.823 | 72 % |
| *Saturnispora dispora*<br>CBS 794 | 0.66 ± 0.00 | 0.828 | 80 % |
| *Phaffomyces*<br>*thermotolerans* CBS 7012 | 0.58 ± 0.01 | 0.824 | 70 % |
| *Cyberlindnera jadinii*<br>CBS 621 | 0.73 ± 0.00 | 0.829 | 88 % |
| *Ogataea parapolyomorpha*<br>CBS 11895 | 0.67 ± 0.01 | 0.835 | 80 % |

Remarkably, the varying macromolecular composition of the yeast species had only little impact on the theoretical maximum yields, which range from 0.823-0.835 g g^-1^ (Table 2). This might indicate that the fraction of substrate dissimilated for the generation of the required amount of ATP *in vivo* has a much bigger impact on yields than the biomass composition itself. The Complex-I-negative yeasts *S. cerevisiae* and *K. lactis* reached 71 and 67%, respectively, of the theoretical maximum biomass yield in chemostat cultures. Similar percentages were calculated for *P. ethanolica* (72%) and *P. thermotolerans* (70%), despite the presence in its genome of genes encoding Complex-I subunits. *S. dispora*, *O. parapolymorpha* (both 80%) and, in particular, *C. jadinii* (88%), more closely approached the theoretical maximum biomass yield.

## Discussion

The overwhelming majority of quantitative yeast physiology studies is based on the use of sugars as carbon- and energy sources. By exploring a small sample of the biodiversity of ascomycete budding yeasts, this study revealed a wide range of growth rates in cultures grown on a synthetic medium with ethanol as sole carbon source. Ethanol-grown cultures of multiple tested strains matched or even exceeded the specific growth rate of approximately 0.4 h^-1^ that is observed for laboratory strains of *S. cerevisiae* when grown on synthetic medium with glucose (van Dijken et al., 2000). An enrichment of high growth rates was observed among all tested strains that belonged to the Phaffomycetales order, although fast growth was observed across the budding yeast subphylum. These results indicate that a broader exploration of intra- and interspecies diversity of yeasts with respect to growth rates on ethanol is highly relevant and may potentially yield even faster-growing strains. Some unexpected phenotypes observed in this study, such as the atypical growth kinetics of the fastest-growing strain *Pichia kudriavzevii* CBS 5147 (Fig. S3, Supporting Information), merit further investigation.

Biomass yields of seven selected yeast strains in aerobic, ethanol-limited chemostat cultures were consistent with the hypothesis that an active Complex-I proton-translocating NADH dehydrogenase is required for achieving high biomass yields on ethanol. However, the low biomass yields of *P. ethanolica* and *P. thermotolerans*, as well as the different biomass yields of *C. jadinii* and *S. dispora* and *O. parapolymorpha* (Fig. 4, Table 1) showed that presence of Complex-I genes does not in itself allow for quantitative prediction of biomass yields. Further research should reveal how regulation of the synthesis of ETS components affects biomass yields in ethanol-grown cultures and which additional levels of regulation are involved in distribution of electrons over proton-coupled and non-proton-coupled branches. Such studies are not only of fundamental interest but can also identify metabolic engineering targets for improving growth energetics in industrial yeasts such as *O. parapolymorpha* (Juergens et al., 2020).

The biomass yield determined for *C. jadinii* (0.73 g g^-1^) in ethanol-limited cultures at a dilution rate of 0.10 h^-1^ is higher than previously reported (Verduyn et al., 1991). This high biomass yield, its high protein content (59 ± 1 wt-%) and its long-term application as ‘ fodder yeast’ in the previous century (Lallemand, 2024; Watteeuw et al., 1979) make it a highly interesting candidate for single-cell protein from ethanol. A recent study in which biomass yields and protein contents of two bacterial and three yeast strains were compared in ethanol-grown shake-flask cultures (van Peteghem et al., 2022b) showed an even higher biomass yield (0.82 g g^-1^) for the related yeast species *Cyberlindnera saturnus* in batch cultures. The chemostat experiments in this study were performed at a relatively low growth rate (0.10 h^-1^). To predict performance of yeasts such as *C. jadinii* and *O. parapolymorpha* in industrial fed-batch cultures, an extended in-depth analysis of growth yield, kinetics and energetics is required. In particular, such studies should cover a range of growth rates that are representative for the dynamic growth-rate regimes in large-scale industrial fed-batch processes.

The observed biomass yield of *C. jadinii* in ethanol-limited chemostat cultures approached the theoretical maximum (88%) at which all ATP requirements for biomass formation is met by respiration of the ‘excess’ reduced cofactors generated in biosynthetic reactions. This raises interesting questions on how ethanol oxidation is coupled to NADH oxidation via Complex I and on how cells allocate cellular resources to enable high fluxes through Complex I which, also in fungi, is a large, multi-subunit protein complex with up to 35 subunits that has to be functionally assembled at and across the mitochondrial inner membrane (Videira, 1998). Such studies are not only relevant for industrial application, but would also provide interesting models for systems biology studies on eukaryotic growth energetics and resource allocation.

Rates and yields are key parameters for any industrial process. Results from this study demonstrate that choice of yeast host cell and the energy coupling strategy in their metabolism impact these parameters significantly. Selecting and improving yeast strains will be highly relevant for optimisation of ethanol-based industrial biotechnology.

## Acknowledgements

We kindly thank Chris Klomp for his help on the protein and RNA content determination protocols, and Niels Tiemersma for his help on the carbohydrate and lipid content determination protocols. Effie Leijten and Erik de Hulster are gratefully acknowledged for their assistance with the chemostat cultivations. Thank you to Marcel van den Broek for his advise on constructing the phylogenetic trees and to Walter van Gulik for his advise on working with the core metabolic model. Dr J.M. Nicaud is kindly acknowledged for providing the *Y. lipolytica* W29 strain. This research is financed by the NWO Stevin Prize awarded to Jack T. Pronk by the Dutch Research Foundation.

