## Supporting information for "Specific growth rates and growth stoichiometries of Saccharomycotina yeasts on ethanol as sole carbon and energy substrate"

Supplementary Figure 1, 2, 3, 4 and 5

Supplementary Table 1, 2 and 3

Supporting information 1

**Table S1.** Yeast strains investigated in this study. Reasons for not including individual strains in Growth Profiler (GP) experiments and/or in subsequent shake-flask (SF) and chemostat (CH) experiments are indicated in the third column. Semi-quantitative (time to stationary phase, TS) and quantitative (µ_max­_) growth data were determined from GP and SF data, respectively.

| Yeast species | Strain | Cultivation excluded from – reason | TS (h) | µ_max_ (h^-1^) |
| --- | --- | --- | --- | --- |
| *Ambrosiozyma monospora* | CBS 2554 | CH – pseudohyphal growth in SF | 12.4 ± 0.5 |  |
| *Barnettozyma californica* | CBS 252 | CH – irregular growth in lag and log phase in SF | 7.0 ± 0.4 |  |
| *Blastobotrys adeninivorans* | CBS 8244 | SF – time to stationary phase in MTP > 21 h | 35.8 ± 0.5 |  |
| *Cyberlindnera fabianii* | CBS 5481 | CH – long lag phase in SF | 8.1 ± 0.7 | 0.235 ± 0.009 |
| *Cyberlindnera jadinii* | CBS 621 |  | 5.7 ± 0.3 | 0.429 ± 0.003 |
| *Cyberlindnera petersonii* | CBS 5555 | CH – irregular growth in exponential phase in SF | 8.0 ± 0.6 | 0.39 ± 0.03 |
| *Debaryomyces hansenii* | CBS 767 | SF – time to stationary phase in MTP > 21 | 27 ± 2 |  |
| *Dekkera bruxellensis* | CBS 2499 | MTP – no growth in inoculation SF on SM-ethanol |  |  |
| *Eremotecium cymbalariae* | CBS 270.75 | MTP – flocculation in inoculation SF on SM-ethanol |  |  |
| *Hyphopichia burtonii* | CBS 2352 | CH – long lag phase in SF | 11 ± 0.4 | 0.229 ± 0.006 |
| *Kazachstania africana* | CBS 2517 | MTP – no growth in inoculation SF on SM-ethanol |  |  |
| *Kluyveromyces lactis* | CBS 2359 |  | 7.9 ± 0.8 | 0.337 ± 0.001 |
| *Kluyveromyces marxianus* | CBS 6556 | CH - µ_max_ in SF < 0.15 h^-1^ | 16 ± 1 | 0.129 ± 0.002 |
| *Komagataella phaffii* | X-33 | CH - µ_max_ in SF < 0.15 h^-1^ | 20.6 ± 0.9 | 0.124 ± 0.000 |
| *Kuraisha capsulata* | CBS 1993 | SF – time to stationary phase in MTP > 21 h | 30.9 ± 0.5 |  |
| *Lachancea kluyveri* | CBS 3082 | MTP – flocculation in inoculation SF on SM-ethanol |  |  |
| *Lachancea thermotolerans* | CBS 6340 | MTP – flocculation in inoculation SF on SM-ethanol |  |  |
| *Lipomyces oliphaga* | CBS 7107 | SF – time to stationary phase in MTP > 21 h | 23 ± 1 |  |
| *Lipomyces starkeyi* | CBS 2512 | SF – time to stationary phase in MTP > 21 h | 37 ± 2 |  |
| *Lipomyces tetrasporus* | CBS 5910 | SF – time to stationary phase in MTP > 21 h | 29 ± 2 |  |
| *Maudiozyma bulderi* | CBS 8638 | MTP – no growth in inoculation SF on SM-ethanol |  |  |
| *Millerozyma farinosa* | CBS 7064 | CH – irregular growth in lag phase in SF | 12.9 ± 0.9 | 0.27 ± 0.01 |
| *Nakaseomyces delphensis* | CBS 2170 | SF – time to stationary phase in MTP > 21 h | 40 ± 8 |  |
| *Naumovozyma castellii* | CBS 4309 | MTP – no growth in inoculation SF on SM-ethanol |  |  |
| *Ogataea henricii* | CBS 5765 | SF – time to stationary phase in MTP > 21 h | 54 ± 7 |  |
| *Ogataea methanolica* | CBS 6515 | SF – time to stationary phase in MTP > 21 h | 27 ± 6 |  |
| *Ogataea methylivora* | CBS 7300 | SF – no growth in MTP after 196 h |  |  |
| *Ogataea parapolymorpha* | CBS 11895 |  | 8.9 ± 0.9 | 0.301 ± 0.000 |
| *Ogataea polymorpha* | CBS 4732 |  | 13.4 ± 0.5 | 0.295 ± 0.006 |
| *Phaffomyces thermotolerans* | CBS 7012 |  | 7.5 ± 0.4 | 0.400 ± 0.001 |
| *Pichia ethanolica* | CBS 8084 |  | 8.5 ± 0.4 | 0.279 ± 0.001 |
| *Pichia kudriavzevii* | CBS 5147 | CH – irregular growth in lag phase in SF | 10.9 ± 0.9 | 0.462 ± 0.007 |
| *Priceomyces haplophilus* | CBS 2028 | MTP – no growth in inoculation SF on SM-ethanol |  |  |
| *Saccharomyces arboricola* | CBS 10644 | MTP – no growth in inoculation SF on SM-ethanol |  |  |
| *Saccharomyces cerevisiae* | CBS 8066 |  | 13.3 ± 0.9 |  |
| *Saccharomyces cerevisiae* | CEN.PK113-7D |  | 17 ± 1 | 0.158 ± 0.001 |
| *Saccharomyces kudriavzevii* | CBS 8840 | MTP – no growth in inoculation SF on SM-ethanol |  |  |
| *Saccharomyces mikatae* | CBS 8839 | SF – time to stationary phase in MTP > 21 h | 65 ± 4 |  |
| *Saccharomyces paradoxus* | CBS 1190 | SF – time to stationary phase in MTP > 21 h | 40 ± 3 |  |
| *Saturnispora dispora* | CBS 794 |  | 10.3 ± 0.9 | 0.348 ± 0.000 |
| *Scheffersomyces stipitis* | CBS 6054 | SF – time to stationary phase in MTP > 21 h | 22 ± 2 |  |
| *Spathaspora passalidarum* | CBS 10155 | SF – time to stationary phase in MTP > 21 h | 24.4 ± 0.5 |  |
| *Starmera quercuum* | CBS 2283 |  | 11 ± 1 | 0.272 ± 0.007 |
| *Tetrapisispora phaffii* | CBS 4417 | SF – no growth in MTP after 196 h |  |  |
| *Torulaspora delbrueckii* | CBS 813 | SF – time to stationary phase in MTP > 21 h | 55 ± 10 |  |
| *Trigonopsis variabilis* | CBS 1040 | SF – time to stationary phase in MTP > 21 h | 29.9 ± 0.4 |  |
| *Vanderwaltozyma polyspora* | CBS 2163 | SF – no growth in MTP after 196 h |  |  |
| *Wickerhamomyces alni* | CBS 6986 | CH - µ_max_ in SF < 0.15 h^-1^ | 11.7 ± 0.4 | 0.14 ± 0.01 |
| *Wickerhamomyces ciferii* | CBS 111 |  | 9.9 ± 0.6 | 0.374 ± 0.002 |
| *Yarrowia lipolytica* | W29 | CH – long lag phase in SF | 18.8 ± 0.9 | 0.20 ± 0.01 |
| *Zygosaccharomyces rouxii* | CBS 732 | SF – no growth in MTP after 196 h |  |  |
| *Zygotorulaspora mrakii* | CBS 4218 | MTP – no growth in inoculation SF on SM-ethanol |  |  |

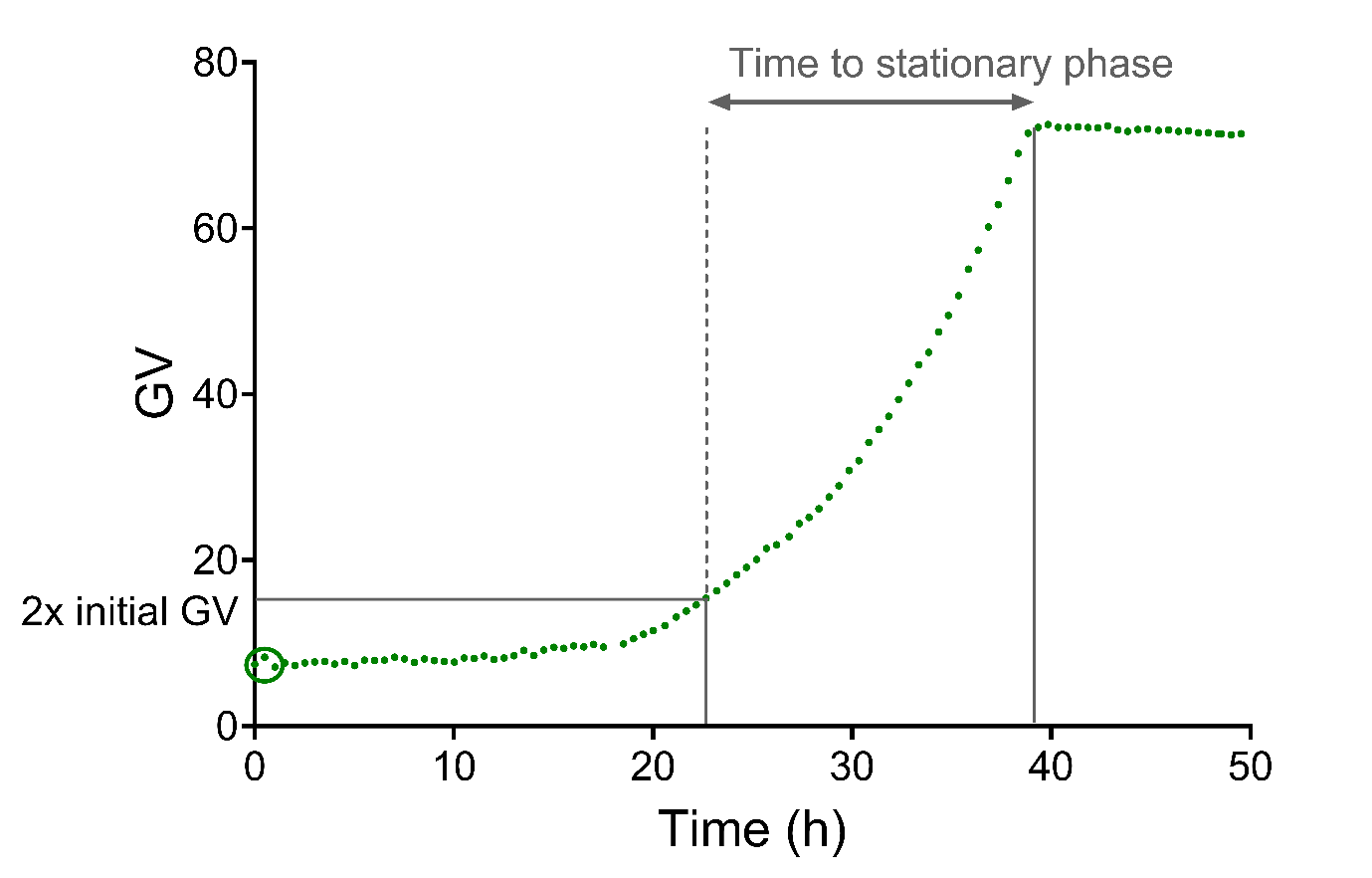

**Figure S1.** Determination of the time to stationary phase from the Growth Profiler cultivation data. The figure shows a representative output for a curve of *Saccharomyces cerevisiae* CEN.PK113-7D (Fig. S2, Supporting Information). The green value (GV, i.e. read-out of the image-analysis-based system) plotted on the y-axis is corrected for the relative position of the well on the microtiter plate. The initial GV was calculated from the first 3 time points (30 min intervals, green circle). To correct for different lag phases of yeast strains, the time point at which GV had doubled from its initial value (start grey double-pointed arrow) is taken as reference (here 22.67 h), at which time GV had doubled from 7.67 to 15.2. The inflection point was reached at 39.34h, which yielded a ‘time to stationary phase’ of 39.34 – 22.67 = 16.67 h.

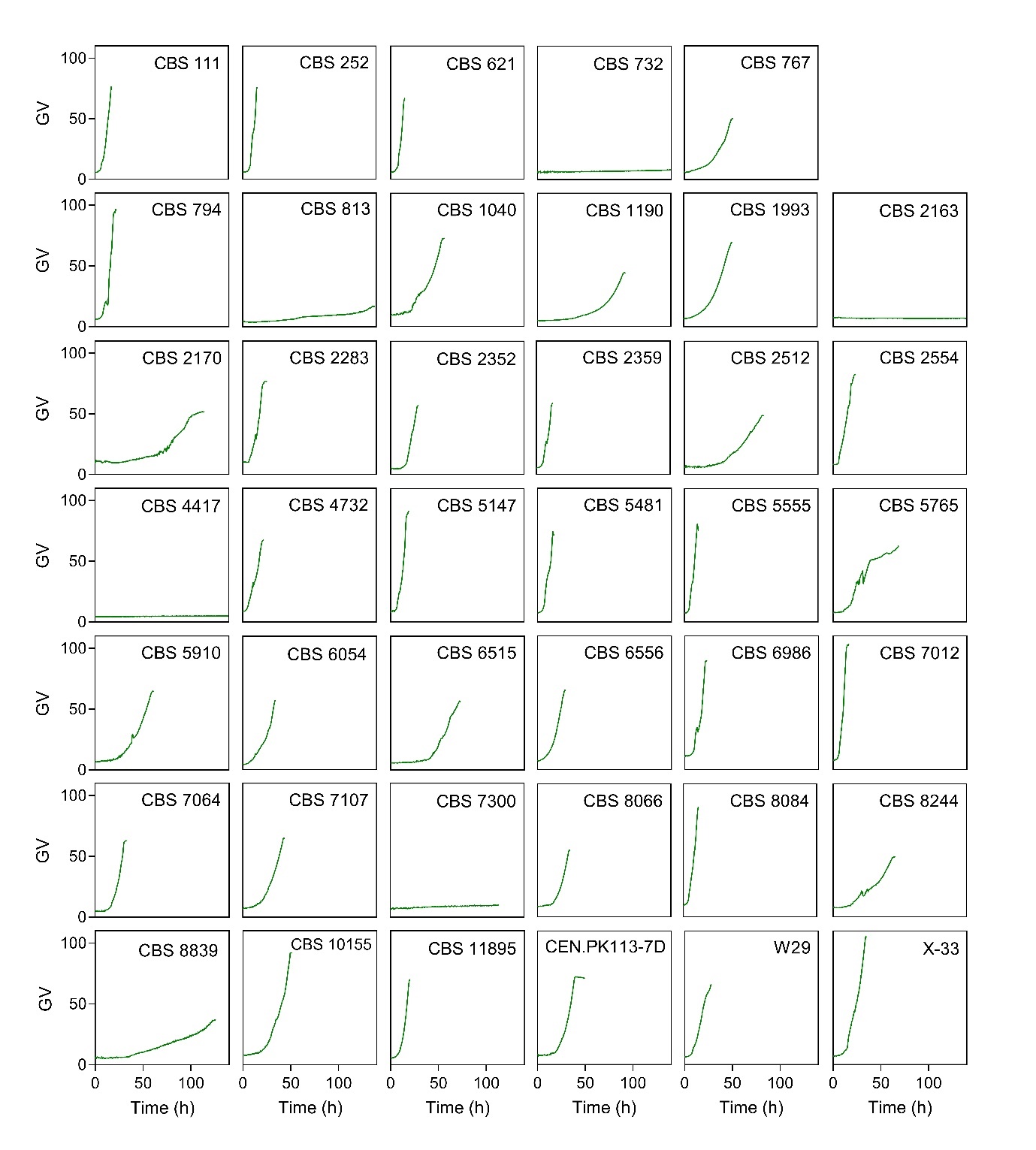

**Figure S2.** Growth curves of strains of 41 yeast species characterised in microtiter plates (n > 4 for each strain, single representative growth curves are shown in the figure). The green value (GV) plotted on the y-axis is corrected for the relative position of the well on the microtiter plate. Green values have a non-linear and strain-dependent correlation to biomass concentration. Each curve is plotted until the onset of the stationary phase. ‘Time to stationary phase’ was calculated as shown in Fig. S1 (Supporting Information).

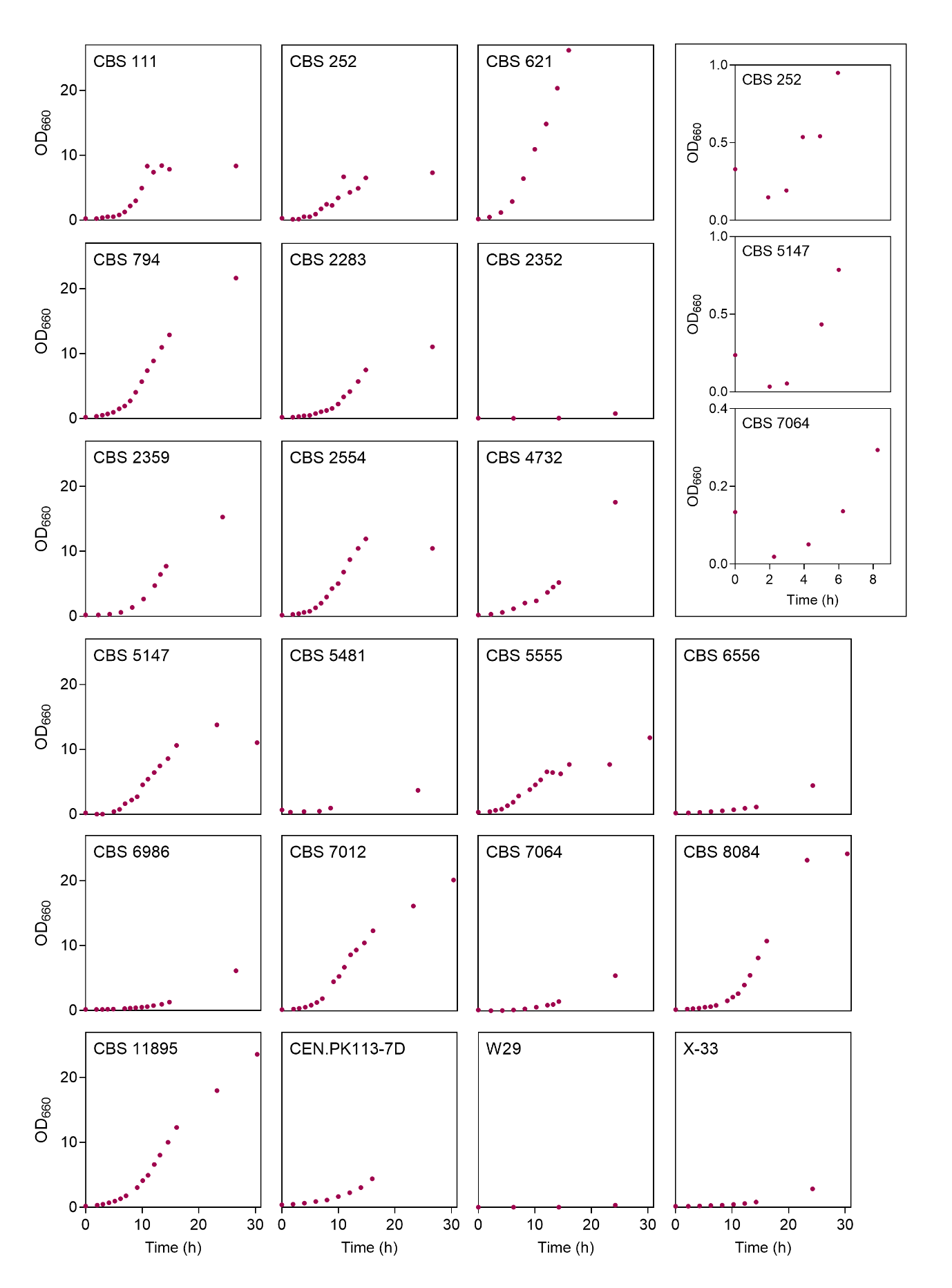

**Figure S3.** Growth curves of strains of 21 yeast species characterised in shake flasks (n > 2 for each strain, single representative curves shown here in the figure. Specific growth rates (µ_max_, Fig. 2) of shake-flask cultures were calculated by linear regression of the relation between ln(OD_660_) and time. For strains considered for chemostat characterisation, at least 6 points over at least 3 biomass doublings in the exponential phase were used to determine µ_max_. The optical density (OD_660_) of strains *Barnettozyma californica* CBS 252, *Pichia kudriavzevii* CBS 5147, and *Millerozyma farinosa* CBS 7064, visualised in three panels in the top-right box, dropped after inoculation.

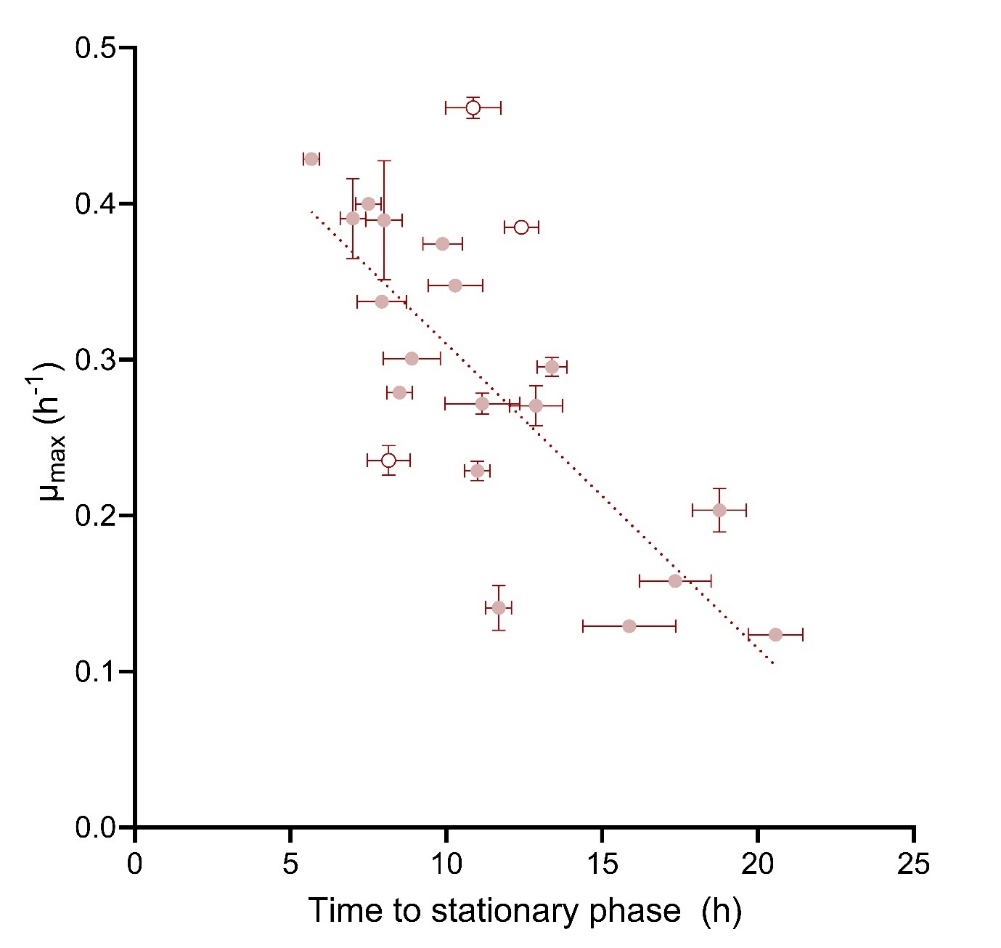

**Figure S4.** Correlation between µ_max_ as calculated from shake-flask cultures and the time to stationary phase as calculated from green values measured in the growth-profiler. Each data point represents a single species. *Pichia kudriavzevii* CBS 5147, *Ambrosiozyma monospora* CBS 2554, *Cyberlindnera fabianii* CBS 5481 (indicated by open symbols from top to bottom) displayed atypical growth characteristics in shake flaks that led to exclusion from chemostat characterisation (Fig. S3 and Table S1, Supporting Information). The correlation coefficient (r^2^) of the linear fit increased from 0.51 to 0.70 (plotted here) upon exclusion of these three species.

**Table S2.** Overview tBLASTn search to establish the electron transport system composition of the Saccharomycotina species considered for chemostat characterisation. The E value with the corresponding percent identity match between brackets, for hits with E value < 0.001 and identity match > 30%, are displayed. Protein sequences of the fungus *Neurospora crassa* were used as queries (columns), except for Complex III subunits where protein sequences from the *S. cerevisiae* CEN.PK strain were retrieved from SGD. The reference sequence from NCBI was used whenever multiple genome datasets were available for a species (rows), the tBLASTn algorithm was run on the NCBI website using default settings.

|  | Control | | Complex I | | |  |  |
| --- | --- | --- | --- | --- | --- | --- | --- |
| Organism (genome data set accession number) | >CAD21159.1 acetyl-CoA synthetase [Neurospora crassa] | >XP_963253.2 ATP synthase beta subunit [Neurospora crassa OR74A] | >XP_957188.3 NADH:ubiquinone oxidoreductase 78 [Neurospora crassa OR74A] | >XP_965490.3 NADH:ubiquinone oxidoreductase 20.9kD subunit [Neurospora crassa OR74A] | >XP_965665.2 NADH:ubiquinone oxidoreductase 49kD subunit [Neurospora crassa OR74A] | >XP_959008.1 alternative NADH-dehydrogenase [Neurospora crassa OR74A] | >EAA32850.1 alternative oxidase [Neurospora crassa OR74A] |
| *Cyberlindnera jadinii*  GCA_001245095.1 | 0.0 (63.12%); 0.0 (61.69%) | 0.0 (84.34%) | 0.0 (65.38%) | 4e-28 (40.00%) | 0.0 (68.92%) | 1e-123 (51.55%);  4e-122 (44.70%) | 4e-69 (40.88%) |
| *Kluyveromyces lactis*  GCF_000002515.2 | 0.0 (61.73%); 0.0 (62.48%) | 0.0 (82.75%) | No significant hit | No significant hit | No significant hit | 3e-141 (51.00%);  2e-114 (41.96%);  1e-64 (33.09%) | No significant hit |
| *Ogataea parapolymorpha*  GCF_000187245.1 | 0.0 (66.87%) | 0.0 (84.79%) | 0.0 (58.01%) | 3e-33 (42.77%) | 0.0 (81.57%) | 4e-149 (46.86%);  1e-138 (47.69%);  3e-59 (36.00%) | No significant hit |
| *Ogataea polymorpha*  GCF_001664045.1 | 0.0 (66.72%) | 0.0 (84.79%) | 0.0 (57.54%) | 5e-33 (42.77%) | 0.0 (81.06%) | 4e-148 (46.47%);  4e-135 (47.89%);  3e-58 (35.38%) | No significant hit |
| *Phaffomyces thermotolerans*  GCA_003707215.2 | 0.0 (61.61%) | 0.0 (83.00%) | 0.0 (63.82%) | 8e-29 (37.20%) | 0.0 (82.47%) | 4e-145 (50.50%);  4e-67 (39.88%) | 5e-63 (43.75%) |
| *Pichia ethanolica*  GCA_030570435.1 | 0.0 (64.85%); 0.0 (59.40%) | 0.0 (82.96%); | 0.0 (57.96%) | 4e-36 (44.24%) | 0.0 (80.56%) | 7e-140 (44.83%);  2e-58 (35.67%) | 2e-69 (43.87%) |
| *Saccharomyces cerevisiae*  GCF_000146045.2 | 0.0 (60.74%);  0.0 (62.28%) | 0.0 (83.15%) | No significant hit | No significant hit | No significant hit | 3e-139 (50.00%);  4e-121 (46.61%) | No significant hit |
| *Saturnispora dispora*  GCA_003243065.1 | 0.0 (65.91%);  0.0 (59.58%) | 0.0 (83.18%) | 0.0 (59.88%) | 3e-33 (42.51%) | 0.0 (81.31%) | 9e-142 (45.91%);  8e-56 (36.31%) | 3e-64 (40.13%) |
| *Starmera quercuum*  GCA_003705275.1 | 0.0 (64.36%);  0.0 (61.36%) | 0.0 (83.41%) | No significant hit | No significant hit | No significant hit | 1e-146 (50.20%);  5e-120 (44.12%);  1e-65 (39.08%) | No significant hit |
| *Wickerhamomyces ciferii*  GCF_000313485.1 | 0.0 (61.70%); 0.0 (62.20%) | 0.0 (84.56%) | 0.0 (67.15%) | 1e-40 (43.45%) | 0.0 (82.22%) | 7e-144 (48.70%);  4e-127 (44.81%);  4e-67 (40.06%) | 2e-67 (47.81%) |

**Table S2** – continued.

|  | Complex III | | |  | Complex IV | |
| --- | --- | --- | --- | --- | --- | --- |
| Organism (genome data set accession number) | SGD:S000005591  ubiquinol--cytochrome-c reductase catalytic subunit CYT1 | SGD:S000000141 ubiquinol--cytochrome-c reductase subunit COR1 | SGD:S000000750  ubiquinol--cytochrome-c reductase catalytic subunit RIP1 | >XP_956486.2 cytochrome c [Neurospora crassa OR74A] | >XP_963448.1 cytochrome c oxidase subunit IV [Neurospora crassa OR74A] | >XP_011394700.1 cytochrome c oxidase polypeptide VI [Neurospora crassa OR74A] |
| *Cyberlindnera jadinii*  GCA_001245095.1 | 7e-151 (76.29%) | 2e-133 (48.37%) | 1e-105 (78.06%) | 7e-46 (64.42%) | 4e-33 (40.94%);  2e-11 (35.71%) | 2e-28 (58.33%) |
| *Kluyveromyces lactis*  GCF_000002515.2 | 4e-157 (78.57%) | 5e-171 (58.70%) | 3e-122 (82.87%) | 1e-46 (67.31%) | 4e-26 (37.42%);  5e-05 (39.22%) | 8e-27 (42.96%) |
| *Ogataea parapolymorpha*  GCF_000187245.1 | 6e-150 (72.55%) | 1e-82 (36.77%) | 2e-96 (68.87%) | 3e-49 (70.19%) | 9e-23 (35.03%) | 2e-28 (43.15%) |
| *Ogataea polymorpha*  GCF_001664045.1 | 6e-150 (72.22%) | 5e-84 (37.22%) | 2e-96 (68.87%) | 2e-49 (69.23%) | 9e-23 (35.03%) | 5e-28 (46.77%) |
| *Phaffomyces thermotolerans*  GCA_003707215.2 | 3e-150 (76.39%) | 1e-143 (49.67%) | 5e-103 (75.51%) | 4e-44 (63.46%) | 2e-33 (44.62%) | 5e-30 (46.31%) |
| *Pichia ethanolica*  GCA_030570435.1 | 1e-142 (73.14%) | 5e-72 (37.36%) | 3e-96 (72.50%) | 7e-47 (64.15%) | 3e-20 (36.72%);  6e-08 (28.87%) | 2e-29 (50.83%) |
| *Saccharomyces cerevisiae*  GCF_000146045.2 | 0.0 (92.43%) | 0.0 (100.00%) | 2e-147 (100.00%) | 1e-48 (69.52%); 1e-48 (69.23%) | 1e-23 (34.42%);  3e-23 (37.18%);  1e-06 (32.50%) | 6e-27 (55.43%) |
| *Saturnispora dispora*  GCA_003243065.1 | 1e-144 (69.28%) | 1e-72 (36.45%) | 2e-99 (70.09%) | 8e-48 (67.62%) | 1e-21 (34.21%);  8-07 (30.38%) | 4e-20 (57.33%) |
| *Starmera quercuum*  GCA_003705275.1 | 2e-149 (74.40%) | 9e-144 (50.66%) | 1e-103 (76.53%) | 1e-46 (66.35%) | 1e-28 (39.22%);  3e-13 (42.17%) | 9e-32 (46.71%) |
| *Wickerhamomyces ciferii*  GCF_000313485.1 | 5e-153 (75.08%) | 9e-139 (50.11%) | 1e-104 (70.05%) | 8e-45 (65.35%) | 6e-28 (40.46%) | 2e-29 (45.27%) |

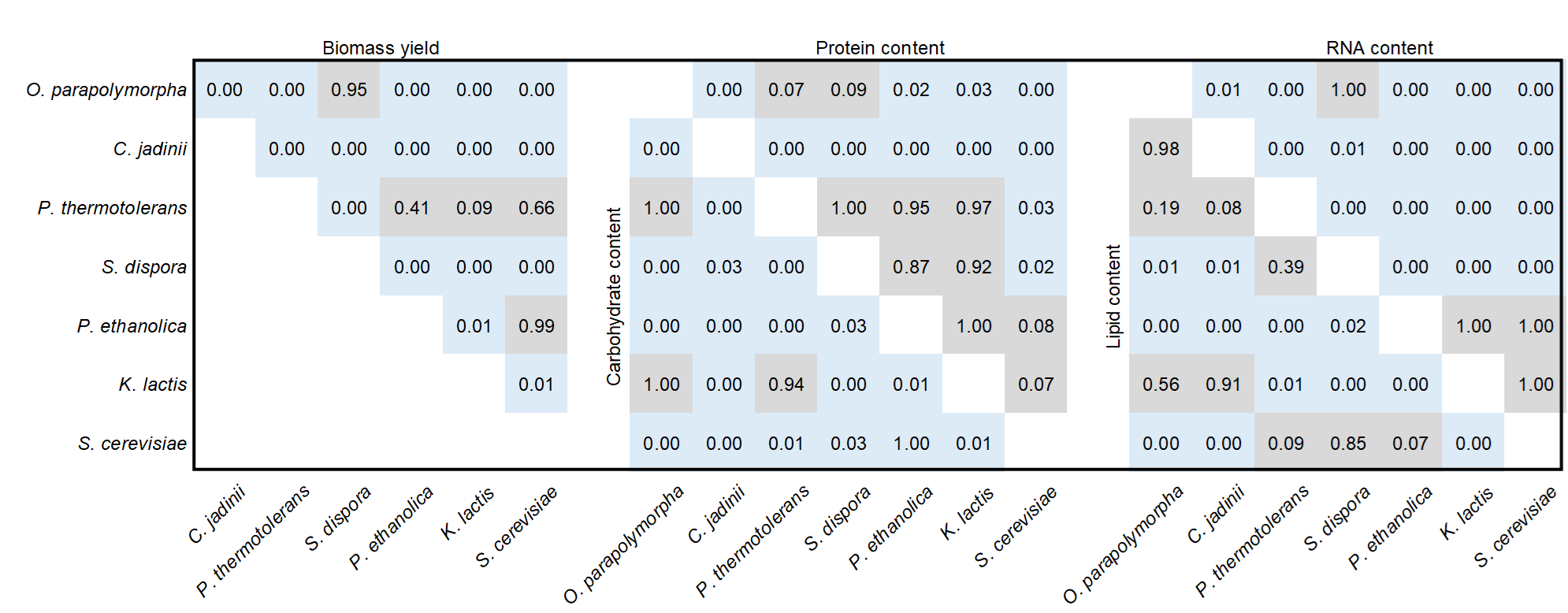

**Figure S5.** Heat map displaying adjusted p values that resulted from statistical analysis of the biomass components analysed for the seven Saccharomycotina species that were grown in ethanol-limited chemostats (30 °C, pH 5.0, dilution rate 0.10 h^-1^). Biomass yield values as g dry weight [g ethanol]^-1^ were compared, and macromolecule content values were compared as g macromolecule [g dry weight]^-1^. Data were analysed using ordinary one-way ANOVA, followed by multiple comparisons where p values were corrected using Tukey’s test.

**Supporting information 1: Stoichiometric growth model used for predicting theoretical maximum biomass yields.**

Ethanol is a more reduced carbon-containing compound than biomass, i.e. ethanol has 6 as degree of reduction per Cmol against 4.2 as degree of reduction per Cmol biomass, which follows from the scenario where ammonium (NH_4_^+^, reference compound set to γ = 0) is used as nitrogen source for biomass formation (Roels, 1980). Therefore, a theoretical scenario exists where ATP requirements for assimilation are completely met by reoxidation of the reducing equivalents generated during assimilation (i.e. NADH and QH_2_) via a respiratory chain coupled to an ATP synthase (Verduyn et al., 1991). In this scenario, the biomass yield is assimilation‑limited and can be determined with a core metabolic network of yeast (Daran-Lapujade et al., 2004).

The biomass formation reaction for each yeast species was based on the experimentally determined protein, RNA, lipid, and carbohydrate content (Table 1). The corresponding molecular weight of 1 Cmol biomass was determined using the generalised molecular formulas of the five macromolecular constituents of biomass, i.e. the components ‘CARBOHYDRATES’, ‘DNA’, ‘LIPID’, PROTEIN’, and ‘RNA’ as listed in the model (see below).

The H^+^/ATP ratio in the ‘mATPase’ reaction of the model (see reaction list below) was minimised from the native ratio, i.e. 10 protons per 3 ATP molecules (Stock et al., 1999), whilst preventing net ATP formation in the mitochondria as a consequence of oxidating reduced cofactors released in assimilatory reactions as a lower boundary.

The model was solved assuming a steady state (S∙v=0), expressing all metabolic fluxes as a function of growth rate and maintenance. The maintenance was set to 0, since this scenario has the underlying assumption that all ATP requirements are met via oxidation of the reduced cofactors released during assimilation.

The theoretical maximum biomass yield was obtained from the model solution by dividing the coefficient of the biomass formation reaction (i.e. 1) by the coefficient of the uptake rate of ethanol from ethanol from the exterior environment.

The P/O ratio was obtained from the solved model by dividing the flux through the ‘mATPase’ reaction (multiplied by 3 because of the reaction stoichiometry in the model) by the sum of the fluxes ‘cNADH deh’, ‘mQH2 deh’, and ‘mNADH deh’. The equations can be found in the reaction list under the category ‘oxidative phosphorylation’.

**Table S3.** H^+^/ATP and P/O ratios required for the predicted biomass yield to become assimilation-limited.

| **Organism** | **H^+^/ATP** | **P/O** |
| --- | --- | --- |
| *S. cerevisiae* | 2.90 | 1.52 |
| *K. lactis* | 3.03 | 1.47 |
| *P. ethanolica* | 2.87 | 1.54 |
| *S. dispora* | 2.87 | 1.54 |
| *P. thermolerans* | 2.97 | 1.50 |
| *O. parapolymorpha* | 3.00 | 1.49 |
| *C. jadinii* | 2.83 | 1.55 |
| Yeast ATP synthase | 3.33 | - |

The reaction list used to generate the core stoichiometric matrix is given below.

```
amino acid synthesis
         alanine transaminase; cALA N-trans
                  1 GLM:cyt + 1 PYR:cyt <=> 1 ALA:cyt + 1 OGL:cyt
         arginine synthesis; cARG syn
                  1 ASP:cyt + 1 ATP:cyt + 1 CARP:cyt + 1 ORN:cyt <=> 1 AMP:cyt + 1 ARG:cyt + 1 FUM:cyt + 3 H:cyt + 1 Pi:cyt + 1 PPi:cyt
         asparagine synthesis; cASN syn
                  1 ASP:cyt + 1 ATP:cyt + 1 GLN:cyt + 1 H2O:cyt <=> 1 ADP:cyt + 1 ASN:cyt + 1 GLM:cyt + 1 H:cyt + 1 Pi:cyt
         asparatate aminotransferase; cASP N-trans
                  1 GLM:cyt + 1 OXACT:cyt <=> 1 ASP:cyt + 1 OGL:cyt
         aspartate kinase; cASP kin
                  1 ASP:cyt + 1 ATP:cyt + 2 H:cyt + 2 NADPH:cyt <=> 1 ADP:cyt + 1 HSER:cyt + 2 NADP:cyt + 1 Pi:cyt
         branched chain amino acid transferase (isoleucine) mit; mILE N-trans
                  1 GLM:mit + 1 H:mit + 1 NADPH:mit + 1 OBU:mit + 1 PYR:mit <=> 1 CO2:mit + 1 H2O:mit + 1 ILE:mit + 1 NADP:mit + 1 OGL:mit
         branched chain amino acid transferase (leucine); cLEU N-trans
                  1 GLM:cyt + 1 IPM:cyt + 1 NAD:cyt <=> 1 CO2:cyt + 1 LEU:cyt + 1 NADH:cyt + 1 OGL:cyt
         branched chain amino acid transferase (valine) cyt; cVAL N-trans
                  1 GLM:cyt + 1 OIV:cyt => 1 OGL:cyt + 1 VAL:cyt
         branched chain amino acid transferase (valine) mit; mVAL N-trans
                  1 GLM:mit + 1 OIV:mit <=> 1 OGL:mit + 1 VAL:mit
         carbamoyl phoshate synthase; cCARP syn
                  2 ATP:cyt + 1 CO2:cyt + 1 GLN:cyt + 2 H2O:cyt <=> 2 ADP:cyt + 1 CARP:cyt + 1 GLM:cyt + 3 H:cyt + 1 Pi:cyt
         cysteine synthese; cCYS syn
                  2 H:cyt + 1 HCYS:cyt + 1 SER:cyt <=> 1 CYS:cyt + 1 NH4:cyt + 1 OBU:cyt
         glutamate ammonia ligase; cGLM N-lig
                  1 ATP:cyt + 1 GLM:cyt + 1 NH4:cyt <=> 1 ADP:cyt + 1 GLN:cyt + 1 H:cyt + 1 Pi:cyt
         glutamate dehydrogenase; cGLM deh
                  1 H:cyt + 1 NADPH:cyt + 1 NH4:cyt + 1 OGL:cyt <=> 1 GLM:cyt + 1 H2O:cyt + 1 NADP:cyt
         glycine hydroxymethyl transferase; cGLY transf
                  1 SER:cyt + 1 THF:cyt <=> 1 GLY:cyt + 1 H2O:cyt + 1 METHF:cyt
         histidine synthesis; cHIS syn
                  1 ATP:cyt + 1 GLN:cyt + 3 H2O:cyt + 2 NAD:cyt + 1 PRPP:cyt <=> 1 AICAR:cyt + 6 H:cyt + 1 HIS:cyt + 2 NADH:cyt + 1 OGL:cyt + 1 Pi:cyt + 2 PPi:cyt
         homocitrate synthesis; cHCIT syn
                  1 ACCoA:cyt + 1 H2O:cyt + 1 OGL:cyt <=> 1 CoA:cyt + 1 H:cyt + 1 HCIT:cyt
         homocysteine synthesis; cHCYS syn
                  1 ACCoA:cyt + 1 H2S:cyt + 1 HSER:cyt <=> 1 ACT:cyt + 1 CoA:cyt + 2 H:cyt + 1 HCYS:cyt
         isopropylmalate synthase cyt; cIPM syn
                  1 ACCoA:cyt + 1 H2O:cyt + 1 OIV:cyt => 1 CoA:cyt + 1 H:cyt + 1 IPM:cyt
         methionine synthase; cMET syn
                  1 H:cyt + 1 HCYS:cyt + 1 MYTHF:cyt <=> 1 MET:cyt + 1 THF:cyt
         ornithine synthesis; mORN syn
                  1 ATP:mit + 2 GLM:mit + 1 H:mit + 1 NADPH:mit => 1 ADP:mit + 1 NADP:mit + 1 OGL:mit + 1 ORN:mit + 1 Pi:mit
         oxoadipate synthesis; mOAD syn
                  1 HCIT:mit + 1 NAD:mit <=> 1 CO2:mit + 1 NADH:mit + 1 OAD:mit
         oxoisovalerate synthesis mit; mOIV syn
                  2 H:mit + 1 NADPH:mit + 2 PYR:mit <=> 1 CO2:mit + 1 H2O:mit + 1 NADP:mit + 1 OIV:mit
         phenylalanine synthesis; cPHE syn
                  1 CHO:cyt + 1 GLM:cyt + 1 H:cyt <=> 1 CO2:cyt + 1 H2O:cyt + 1 OGL:cyt + 1 PHE:cyt
         proline dehydrogenase; cPRO deh
                  1 ATP:cyt + 1 GLM:cyt + 2 H:cyt + 2 NADPH:cyt <=> 1 ADP:cyt + 1 H2O:cyt + 2 NADP:cyt + 1 Pi:cyt + 1 PRO:cyt
         serine synthesis; cSER syn
                  1 3PG:cyt + 1 GLM:cyt + 1 H2O:cyt + 1 NAD:cyt <=> 1 H:cyt + 1 NADH:cyt + 1 OGL:cyt + 1 Pi:cyt + 1 SER:cyt
         shikimate pathway; cSHI path
                  1 ATP:cyt + 1 E4P:cyt + 1 NADPH:cyt + 2 PEP:cyt <=> 1 ADP:cyt + 1 CHO:cyt + 1 NADP:cyt + 4 Pi:cyt
         threonine aldolase; cTHR ald
                  1 THR:cyt => 1 ACTAL:cyt + 1 GLY:cyt
         threonine dehydratase mit; mTHR deh
                  1 H:mit + 1 THR:mit => 1 NH4:mit + 1 OBU:mit
         threonine synthesis; cTHR syn
                  1 ATP:cyt + 1 H2O:cyt + 1 HSER:cyt <=> 1 ADP:cyt + 1 H:cyt + 1 Pi:cyt + 1 THR:cyt
         thryptophan synthesis; cTRP syn
                  1 CHO:cyt + 1 GLN:cyt + 1 PRPP:cyt + 1 SER:cyt <=> 1 CO2:cyt + 1 GAP:cyt + 1 GLM:cyt + 1 H:cyt + 1 H2O:cyt + 2 Pi:cyt + 1 PYR:cyt + 1 TRP:cyt
         tyrosine synthesis; cTYR syn
                  1 CHO:cyt + 1 GLM:cyt + 1 NADP:cyt <=> 1 CO2:cyt + 1 NADPH:cyt + 1 OGL:cyt + 1 TYR:cyt

biomass formation
biomass formation C. jadinii; biom-Cj
                  0,198 CARBHYD:cyt + 0,00380 DNA:cyt + 0,0987 LIPID:cyt + 0,625 PROT:cyt + 0,0749 RNA:cyt => 1 biom-Cj:ext
biomass formation K. lactis; biom-Kl
                  0,322 CARBHYD:cyt + 0,00403 DNA:cyt + 0,0990 LIPID:cyt + 0,523 PROT:cyt + 0,0522 RNA:cyt => 1 biom-Kl:ext
biomass formation O. parapolymorpha; biom-Op
                  0,299 CARBHYD:cyt + 0,00372 DNA:cyt + 0,0994 LIPID:cyt + 0,533 PROT:cyt + 0,0651 RNA:cyt => 1 biom-Op:ext
biomass formation P. thermotolerans; biom-Pt
                  0,321 CARBHYD:cyt + 0,00395 DNA:cyt + 0,120 LIPID:cyt + 0,523 PROT:cyt + 0,0331 RNA:cyt => 1 biom-Pt:ext
biomass formation P. ethanolica; biom-Pe
                  0,272 CARBHYD:cyt + 0,00397 DNA:cyt + 0,155 LIPID:cyt + 0,516 PROT:cyt + 0,0509 RNA:cyt => 1 biom-Pe:ext
biomass formation S. cerevisiae; biom-Sc
                  0,291 CARBHYD:cyt + 0,00424 DNA:cyt + 0,147 LIPID:cyt + 0,503 PROT:cyt + 0,0543 RNA:cyt => 1 biom-Sc:ext
Biomass formation S. dispora; biom-Sd
                  0,246 CARBHYD:cyt + 0,00408 DNA:cyt + 0,135 LIPID:cyt + 0,546 PROT:cyt + 0,0707 RNA:cyt => 1 biom-Sd:ext

C-1 metabolism
         dihydrofolate reductase; cDHF red
                  1 DHF:cyt + 1 H:cyt + 1 NADPH:cyt => 1 NADP:cyt + 1 THF:cyt
         methylenetetrahydrofolate dehydrogenase; cMETHF deh
                  1 FTHF:cyt + 1 H:cyt + 1 NADPH:cyt => 1 H2O:cyt + 1 METHF:cyt + 1 NADP:cyt
         methylenetetrahydrofolate reductase; cMETHF red
                  1 H:cyt + 1 METHF:cyt + 1 NADPH:cyt => 1 MYTHF:cyt + 1 NADP:cyt

catabolism
         sulfate assimilation; cSO4 ass
                  2 ATP:cyt + 3 H:cyt + 4 NADPH:cyt + 1 SO4:cyt => 1 ADP:cyt + 1 AMP:cyt + 2 H2O:cyt + 1 H2S:cyt + 4 NADP:cyt + 1 Pi:cyt + 1 PPi:cyt

diffusion
         CO2 diffusion; eCO2<-cCO2
                  1 CO2:cyt <=> 1 CO2:ext
         intracellular carbondioxide diffusion; cCO2<-mCO2
                  1 CO2:mit => 1 CO2:cyt
         intracellular oxygen diffusion; cO2->mO2
                  1 O2:cyt => 1 O2:mit
         intracellular water diffusion; cH2O<-mH2O
                  1 H2O:mit => 1 H2O:cyt
         oxygen diffusion; eO2->cO2
                  1 O2:ext <=> 1 O2:cyt
         water diffusion; eH2O<-cH2O
                  1 H2O:cyt <=> 1 H2O:ext

gluconeogenesis
         fructose-diphosphatase; cF16P phos
                  1 F16P:cyt + 1 H2O:cyt => 1 F6P:cyt + 1 Pi:cyt
         isocitrate lyase; cICIT lya
                  1 ICIT:cyt => 1 GLYO:cyt + 1 SUC:cyt
         malate synthase; cMAL syn
                  1 ACCoA:cyt + 1 GLYO:cyt + 1 H2O:cyt => 1 CoA:cyt + 1 H:cyt + 1 MAL:cyt
         PEP carboxykinase; cPEP ck
                  1 ATP:cyt + 1 OXACT:cyt => 1 ADP:cyt + 1 CO2:cyt + 1 PEP:cyt

glycolysis, lower
         enolase; cEnol
                  1 2PG:cyt <=> 1 H2O:cyt + 1 PEP:cyt
         glyceraldehyde phosphate dehydrogenase; cGAP deh
                  1 GAP:cyt + 1 NAD:cyt + 1 Pi:cyt <=> 1 13PG:cyt + 1 H:cyt + 1 NADH:cyt
         phosphoglycerate kinase; c13PG kin
                  1 13PG:cyt + 1 ADP:cyt <=> 1 3PG:cyt + 1 ATP:cyt
         phosphoglycerate mutase; c3PG mut
                  1 3PG:cyt <=> 1 2PG:cyt
         pyruvate kinase; cPYR kin
                  1 ADP:cyt + 1 H:cyt + 1 PEP:cyt => 1 ATP:cyt + 1 PYR:cyt

glycolysis, upper
         fructosebisphosphate aldolase; cF16P ald
                  1 F16P:cyt <=> 1 DHAP:cyt + 1 GAP:cyt
         glucose 6-phosphate isomerase; cG6P iso
                  1 G6P:cyt <=> 1 F6P:cyt
         triose phophate isomerase; cTP iso
                  1 DHAP:cyt <=> 1 GAP:cyt

intracellular transport
         ADP-ATP antiport cyt/mit; ADPc<->ATPm
                  1 ADP:cyt + 1 ATP:mit => 1 ADP:mit + 1 ATP:cyt
         mit. ammonium carrier protein; NH4m car
                  1 H:cyt + 1 NH4:mit => 1 H:mit + 1 NH4:cyt
         mit. citrate/oxaloacetate carrier; CTP1m car
                  1 CIT:cyt + 1 H:cyt + 1 OXACT:mit => 1 CIT:mit + 1 H:mit + 1 OXACT:cyt
         mit. homocitrate carrier; HCITm car
                  4 H:cyt + 1 HCIT:cyt => 4 H:mit + 1 HCIT:mit
         mit. isocitrate/oxaloacetate carrier; CTP2m car
                  1 H:mit + 1 ICIT:mit + 1 OXACT:cyt => 1 H:cyt + 1 ICIT:cyt + 1 OXACT:mit
         mit. isoleucine carrier; ILEm car
                  1 H:cyt + 1 ILE:mit => 1 H:mit + 1 ILE:cyt
         mit. ornithine carrier; ORNm car
                  1 H:cyt + 1 ORN:mit => 1 H:mit + 1 ORN:cyt
         mit. oxaloacetate exporter; OXACTm ex
                  1 H:mit + 1 OXACT:mit => 1 H:cyt + 1 OXACT:cyt
         mit. oxobutyrate carrier; OBUm car
                  1 H:cyt + 1 OBU:cyt => 1 H:mit + 1 OBU:mit
         mit. oxogluterate/malate carrier; ODC1m car
                  1 MAL:cyt + 1 OGL:mit => 1 MAL:mit + 1 OGL:cyt
         mit. oxogluterate/oxoadipate carrier; ODC2m car
                  1 OAD:mit + 1 OGL:cyt => 1 OAD:cyt + 1 OGL:mit
         mit. oxoisovalerate carrier; OIVm car
                  1 OIV:mit => 1 OIV:cyt
         mit. phosphate carrier mit; Pi_m car
                  2 H:cyt + 1 Pi:cyt => 2 H:mit + 1 Pi:mit
         mit. pyruvate proton symport; PYRm car
                  2 H:cyt + 1 PYR:cyt => 2 H:mit + 1 PYR:mit
         mit. succinate/fumerate carrier; SFC1m car
                  1 FUM:mit + 1 SUC:cyt => 1 FUM:cyt + 1 SUC:mit
         mit. threonine carrier; THRm car
                  1 H:cyt + 1 THR:cyt => 1 H:mit + 1 THR:mit
         mit. valine importer; VALm imp
                  1 H:cyt + 1 VAL:cyt => 1 H:mit + 1 VAL:mit

lipid synthesis
         acetyl-CoA carboxylase; cACCoA carb
                  1 ACCoA:cyt + 1 ATP:cyt + 1 CO2:cyt + 1 H2O:cyt => 1 ADP:cyt + 2 H:cyt + 1 MACoA:cyt + 1 Pi:cyt
         adenosly homocysteinase; cSAH hyd
                  1 H2O:cyt + 1 SAH:cyt => 1 A:cyt + 1 H:cyt + 1 HCYS:cyt
         average fatty acid formation; cAvFA form
                  1,7 OLE-CoA:cyt + 4,4 PLLM-CoA:cyt + 1,4 PLM-CoA:cyt + 1 STE-CoA:cyt => 8,5 avFA-CoA:cyt
         average phospholipid formation; cAvPL form
                  11 PHD-CHO:cyt + 4 PHD-ETA:cyt + 3 PHD-SER:cyt => 18 avPL:cyt
         FAT formation; cFAT form
                  1 avFA-CoA:cyt + 1 H2O:cyt + 1 PHD:cyt => 1 CoA:cyt + 1 FAT:cyt + 1 Pi:cyt
         glycerol 3-phosphate acyltransferase; cGOH3P trans
                  2 avFA-CoA:cyt + 1 GOH3P:cyt => 2 CoA:cyt + 1 PHD:cyt
         glycerol 3-phosphate dehydrogenase; cGOH3P deh
                  1 DHAP:cyt + 1 H:cyt + 1 NADH:cyt => 1 GOH3P:cyt + 1 NAD:cyt
         methionine adenosyl transferase; cMET Atrans
                  1 ATP:cyt + 2 H2O:cyt + 1 MET:cyt => 1 H:cyt + 3 Pi:cyt + 1 SAM:cyt
         palmitate CoA ligase; cPLM lig
                  1 ATP:cyt + 1 CoA:cyt + 1 H2O:cyt + 1 PLM:cyt => 1 AMP:cyt + 1 H:cyt + 2 Pi:cyt + 1 PLM-CoA:cyt
         palmitate-CoA desaturase; cPLM desat
                  1 H:cyt + 1 NADPH:cyt + 1 O2:cyt + 1 PLM-CoA:cyt => 2 H2O:cyt + 1 NADP:cyt + 1 PLLM-CoA:cyt
         palmitic acid synthesis; cPLM syn
                  1 ACCoA:cyt + 20 H:cyt + 7 MACoA:cyt + 14 NADPH:cyt => 7 CO2:cyt + 8 CoA:cyt + 6 H2O:cyt + 14 NADP:cyt + 1 PLM:cyt
         phosphatidate cytidyl transferase; cPHD-C trans
                  1 CTP:cyt + 1 H2O:cyt + 1 PHD:cyt => 1 CMP-DGOH:cyt + 2 Pi:cyt
         phosphatidyl-ethanolamine methyltransferase; cPHD-EA mtrans
                  1 PHD-ETA:cyt + 3 SAM:cyt => 3 H:cyt + 1 PHD-CHO:cyt + 3 SAH:cyt
         phosphatidyl-serine decarboxylase; cPHD-SER dcarb
                  1 PHD-SER:cyt => 1 CO2:cyt + 1 PHD-ETA:cyt
         phosphatidyl-serine synthase; cPHD-SER syn
                  1 CMP-DGOH:cyt + 1 SER:cyt => 1 CMP:cyt + 1 PHD-SER:cyt
         stearate CoA desaturase; cSTE desat
                  1 H:cyt + 1 NADPH:cyt + 1 O2:cyt + 1 STE-CoA:cyt => 2 H2O:cyt + 1 NADP:cyt + 1 OLE-CoA:cyt
         stearate CoA ligase; cSTE ligase
                  1 ATP:cyt + 1 CoA:cyt + 1 H2O:cyt + 1 STE:cyt => 1 AMP:cyt + 1 H:cyt + 2 Pi:cyt + 1 STE-CoA:cyt
         stearic acid synthesis; cSTE syn
                  1 ACCoA:cyt + 23 H:cyt + 8 MACoA:cyt + 16 NADPH:cyt => 8 CO2:cyt + 9 CoA:cyt + 7 H2O:cyt + 16 NADP:cyt + 1 STE:cyt

macromolecule synthesis
         average amino acid formation; cAvAA form
                  0,977 ALA:cyt + 0,386 ARG:cyt + 0,408 ASN:cyt + 0,52 ASP:cyt + 0,0139 CYS:cyt + 1,02 GLM:cyt + 0,526 GLN:cyt + 0,889 GLY:cyt + 0,193 HIS:cyt + 0,589 ILE:cyt + 0,801 LEU:cyt + 0,657 LYS:cyt + 0,114 MET:cyt + 0,0238 ORN:cyt + 0,376 PHE:cyt + 0,422 PRO:cyt + 0,533 SER:cyt + 0,557 THR:cyt + 0,0649 TRP:cyt + 0,196 TYR:cyt + 0,733 VAL:cyt <=> 10 avAA:cyt
         carbohydrate synthesis; cCARBHYD syn
                  1 ATP:cyt + 1 G6P:cyt + 1 H2O:cyt => 1 ADP:cyt + 6 CARBHYD:cyt + 1 H:cyt + 2 Pi:cyt
         DNA polymerisation; cDNA poly
                  0,3 ATP:cyt + 0,2 CTP:cyt + 0,2 GTP:cyt + 1 H:cyt + 0,3 METHF:cyt + 1 NADPH:cyt + 0,3 UTP:cyt => 0,3 DHF:cyt + 9,8 DNA:cyt + 1 H2O:cyt + 1 NADP:cyt + 1 PPi:cyt
         lipid formation; cLipid form
                  0,45 avPL:cyt + 0,55 FAT:cyt => 47,2 LIPID:cyt
         lysine synthesis; cLYS syn
                  1 ATP:cyt + 2 GLM:cyt + 1 NAD:cyt + 2 NADPH:cyt + 1 OAD:cyt <=> 1 AMP:cyt + 1 LYS:cyt + 1 NADH:cyt + 2 NADP:cyt + 2 OGL:cyt + 1 PPi:cyt
         protein polymerisation; cPROT poly
                  3 ATP:cyt + 1 avAA:cyt + 2 H2O:cyt <=> 2 ADP:cyt + 1 AMP:cyt + 4 H:cyt + 2 Pi:cyt + 1 PPi:cyt + 4,81 PROT:cyt
         RNA polymerisation; cRNA syn
                  0,3 ATP:cyt + 0,2 CTP:cyt + 0,2 GTP:cyt + 0,3 UTP:cyt => 1 PPi:cyt + 9,5 RNA:cyt

maintenance
         maintenance; cMaintenance
                  1 ATP:cyt + 1 H2O:cyt => 1 ADP:cyt + 1 H:cyt + 1 Pi:cyt

nucleotide synthesis
         adenosine kinase; cA kin
                  1 A:cyt + 1 ATP:cyt <=> 1 ADP:cyt + 1 AMP:cyt + 1 H:cyt
         adenylate kinase; cAMP kin
                  1 AMP:cyt + 1 ATP:cyt <=> 2 ADP:cyt
         AMP synthesis; cAMP syn
                  1 ASP:cyt + 1 ATP:cyt + 1 IMP:cyt <=> 1 ADP:cyt + 1 AMP:cyt + 1 FUM:cyt + 2 H:cyt + 1 Pi:cyt
         CTP synthetase; cCTP syn
                  1 ATP:cyt + 1 GLN:cyt + 1 H2O:cyt + 1 UTP:cyt <=> 1 ADP:cyt + 1 CTP:cyt + 1 GLM:cyt + 2 H:cyt + 1 Pi:cyt
         cytidylate kinase; cCMP kin
                  1 ATP:cyt + 1 CMP:cyt <=> 1 ADP:cyt + 1 CDP:cyt
         GMP synthesis; cGMP syn
                  1 ATP:cyt + 1 GLN:cyt + 2 H2O:cyt + 1 IMP:cyt + 1 NAD:cyt <=> 1 AMP:cyt + 1 GLM:cyt + 1 GMP:cyt + 4 H:cyt + 1 NADH:cyt + 1 PPi:cyt
         guanylate kinase; cGMP kin
                  1 ATP:cyt + 1 GMP:cyt => 1 ADP:cyt + 1 GDP:cyt
         IMP synthesis; cIMP syn
                  1 AICAR:cyt + 1 FTHF:cyt => 1 H2O:cyt + 1 IMP:cyt + 1 THF:cyt
         nucleoside diphosphate kinase 1; cGDP kin
                  1 ATP:cyt + 1 GDP:cyt <=> 1 ADP:cyt + 1 GTP:cyt
         nucleoside diphosphate kinase 2; cUDP kin
                  1 ATP:cyt + 1 UDP:cyt => 1 ADP:cyt + 1 UTP:cyt
         nucleoside diphosphate kinase 3; cCDP kin
                  1 ATP:cyt + 1 CDP:cyt => 1 ADP:cyt + 1 CTP:cyt
         phosphoribosyl pyrophosphate synthesis; cPRPP syn
                  1 ATP:cyt + 1 RIBU5P:cyt <=> 1 AMP:cyt + 1 H:cyt + 1 PRPP:cyt
         phosphoribosyl-5-amino 4-imidazole carboxamide; cAICAR syn
                  1 ASP:cyt + 4 ATP:cyt + 1 CO2:cyt + 1 FTHF:cyt + 2 GLN:cyt + 1 GLY:cyt + 2 H2O:cyt + 1 PRPP:cyt <=> 4 ADP:cyt + 1 AICAR:cyt + 1 FUM:cyt + 2 GLM:cyt + 8 H:cyt + 4 Pi:cyt + 1 PPi:cyt + 1 THF:cyt
         pyrophosphatase; cPPi ase
                  1 H2O:cyt + 1 PPi:cyt => 2 Pi:cyt
         UMP synthesis; cUMP syn
                  1 ASP:cyt + 1 CARP:cyt + 0,5 O2:cyt + 1 PRPP:cyt <=> 1 CO2:cyt + 2 H2O:cyt + 1 Pi:cyt + 1 PPi:cyt + 1 UMP:cyt
         uridylate kinase; cUMP kin
                  1 ATP:cyt + 1 UMP:cyt <=> 1 ADP:cyt + 1 UDP:cyt

oxidative phosphorylation
         F1-F0 ATPase; mATPase_wt
                  3 ADP:mit + 10 H:cyt + 3 Pi:mit <=> 3 ATP:mit + 7 H:mit + 3 H2O:mit
         ATPase assimilation-limited scenario C. jadinii; mATPase_Cj_theor
                  3 ADP:mit + 8,5 H:cyt + 3 Pi:mit <=> 3 ATP:mit + 5,5 H:mit + 3 H2O:mit
         ATPase assimilation-limited scenario K. lactis; mATPase_Kl_theor
                  3 ADP:mit + 9,1 H:cyt + 3 Pi:mit <=> 3 ATP:mit + 6,1 H:mit + 3 H2O:mit
         ATPase assimilation-limited scenario O. parapolymorpha; mATPase_Op_theor
                  3 ADP:mit + 9,0 H:cyt + 3 Pi:mit <=> 3 ATP:mit + 6,0 H:mit + 3 H2O:mit
         ATPase assimilation-limited scenario P. thermotolerans; mATPase_Pt_theor
                  3 ADP:mit + 8,9 H:cyt + 3 Pi:mit <=> 3 ATP:mit + 5,9 H:mit + 3 H2O:mit
         ATPase assimilation-limited scenario P. ethanolica; mATPase_Pe_theor
                  3 ADP:mit + 8,6 H:cyt + 3 Pi:mit <=> 3 ATP:mit + 5,6 H:mit + 3 H2O:mit
         ATPase assimilation-limited scenario S. cerevisiae; mATPase_Sc_theor
                  3 ADP:mit + 8,7 H:cyt + 3 Pi:mit <=> 3 ATP:mit + 5,7 H:mit + 3 H2O:mit
         ATPase assimilation-limited scenario S. dispora; mATPase_Sd_theor
                  3 ADP:mit + 8,6 H:cyt + 3 Pi:mit <=> 3 ATP:mit + 5,6 H:mit + 3 H2O:mit

         NADH dehydrogenase cyt; cNADH deh
                  6 H:mit + 1 NADH:cyt + 0,5 O2:mit => 5 H:cyt + 1 H2O:mit + 1 NAD:cyt
         NADH dehydrogenase Complex I mit; mNADH deh CI
                  11 H:mit + 1 NADH:mit + 0,5 O2:mit => 10 H:cyt + 1 H2O:mit + 1 NAD:mit
         NADH dehydrogenase Type II mit; mNADH deh T2
                  7 H:mit + 1 NADH:mit + 0,5 O2:mit => 6 H:cyt + 1 H2O:mit + 1 NAD:mit
         quinol dehydrogenase mit; mQH2 deh
                  1 QH2:mit + 6 H:mit + 0,5 O2:mit => 1 Q:mit + 6 H:cyt + 1 H2O:mit

pentose phosphate pathway
         ribosephosphate isomerase; cRIBU iso
                  1 RIBU5P:cyt <=> 1 RIB5P:cyt
         ribulosephosphate 3-epimerase; cRIBUP epi
                  1 RIBU5P:cyt <=> 1 XYL5P:cyt
         transaldolase; cTA1
                  1 GAP:cyt + 1 SED7P:cyt <=> 1 E4P:cyt + 1 F6P:cyt
         transketolase 1; cTK1
                  1 RIB5P:cyt + 1 XYL5P:cyt <=> 1 GAP:cyt + 1 SED7P:cyt
         transketolase 2; cTK2
                  1 E4P:cyt + 1 XYL5P:cyt <=> 1 F6P:cyt + 1 GAP:cyt

pyruvate branchpoint
         acetaldehyde dehydrogenase (NAD); cACTAL deh (NAD)
                  1 ACTAL:cyt + 1 H2O:cyt + 1 NAD:cyt <=> 1 ACT:cyt + 2 H:cyt + 1 NADH:cyt
         acetaldehyde dehydrogenase (NADP); cACTAL deh (NADP)
                  1 ACTAL:cyt + 1 H2O:cyt + 1 NADP:cyt <=> 1 ACT:cyt + 2 H:cyt + 1 NADPH:cyt
         acetyl-CoA synthase; cACCoA syn
                  1 ACT:cyt + 1 ATP:cyt + 1 CoA:cyt + 1 H2O:cyt => 1 ACCoA:cyt + 1 AMP:cyt + 1 H:cyt + 2 Pi:cyt
         alcohol dehydrogenase; cETOH deh
                  1 ACTAL:cyt + 1 H:cyt + 1 NADH:cyt <=> 1 ETOH:cyt + 1 NAD:cyt

TCA cycle
         aconitase 1 mit; mACON 1
                  1 CIT:mit => 1 ACO:mit + 1 H2O:mit
         aconitase 2 mit; mACON 2
                  1 ACO:mit + 1 H2O:mit => 1 ICIT:mit
         citrate synthase; cCIT syn
                  1 ACCoA:cyt + 1 H2O:cyt + 1 OXACT:cyt => 1 CIT:cyt + 1 CoA:cyt + 1 H:cyt
         fumarate hydrase; cFUM hy
                  1 FUM:cyt + 1 H2O:cyt <=> 1 MAL:cyt
         fumarate hydrase mit; mFUM hy
                  1 FUM:mit + 1 H2O:mit <=> 1 MAL:mit
         isocitrate dehydrogenase (NAD) mit; mICIT deh_NAD
                  1 ICIT:mit + 1 NAD:mit => 1 CO2:mit + 1 NADH:mit + 1 OGL:mit
         isocitrate dehydrogenase (NADP) mit; mICIT deh_NADP
                  1 ICIT:mit + 1 NADP:mit => 1 CO2:mit + 1 NADPH:mit + 1 OGL:mit
         malate dehydrogenase; cMAL deh
                  1 MAL:cyt + 1 NAD:cyt <=> 1 H:cyt + 1 NADH:cyt + 1 OXACT:cyt
         malate dehydrogenase mit; mMAL deh
                  1 MAL:mit + 1 NAD:mit <=> 1 H:mit + 1 NADH:mit + 1 OXACT:mit
         oxogluterate dehydrogenase mit; mOGL deh
                  1 CoA:mit + 1 NAD:mit + 1 OGL:mit => 1 CO2:mit + 1 NADH:mit + 1 SUCCOA:mit
         succinate dehydrogenase mit; mSUC deh
                  1 Q:mit + 1 SUC:mit <=> 1 QH2:mit + 1 FUM:mit
         succinyl CoA synthetase mit; mSUCCoA syn
                  1 ADP:mit + 1 Pi:mit + 1 SUCCOA:mit <=> 1 ATP:mit + 1 CoA:mit + 1 SUC:mit

transport
         ATPase plasmamembrane; eATPase
                  1 ATP:cyt + 1 H2O:cyt => 1 ADP:cyt + 1 H:ext + 1 Pi:cyt
         ethanol transport; eETOH trans
                  1 ETOH:cyt => 1 ETOH:ext
         metal import; eMET imp
                  1 metal:ext => 1 metal:cyt
         NH4 transport; eNH4 trans
                  1 NH4:ext => 1 NH4:cyt
         phosphate transport; ePi trans
                  2 H:ext + 1 Pi:ext => 2 H:cyt + 1 Pi:cyt
         SO4 transport; eSO4 trans
                  3 H:ext + 1 SO4:ext => 3 H:cyt + 1 SO4:cyt

Component List

Name                                         ShortName         Composition
1,3-phosphoglycerate                         13PG               C3H4O10P2-4
2-phosphoglycerate                           2PG                C3H4O7P-3
3-phosphoglycerate                           3PG                C3H4O7P-3
acetaldehyde                                 ACTAL              C2H4O
acetate                                      ACT                C2H3O2-1
acetyl-CoA                                   ACCoA              C23H34N7O17P3S-4
adenosine                                    A                  C10H13N5O4
ADP                                          ADP                C10H12N5O10P2-3
alanine                                      ALA                C3H7NO2
AMP                                          AMP                C10H12N5O7P-2
arginine                                     ARG                C6H15N4O2+1
asparagine                                   ASN                C4H8N2O3
aspartate                                    ASP                C4H6NO4-1
ATP                                          ATP                C10H12N5O13P3-4
average amino acid                           avAA               C4,81H9,61N1,32O2,53S0,0128-0,0282
AVERAGE FATTY ACID-CoA                       avFA-CoA           C37,6H61,8N7O17P3S-4
average phospholipid                         avPL               C40,3H77,3NO8,33P
Biomass C. jadinii                           biom-Cj          CH1,60N0,203O0,436P0,00920S0,00166
Biomass K. lactis                            biom-Kl          CH1,62N0,167O0,490P0,00683S0,00139
Biomass O. parapolymorpha                    biom-Op          CH1,61N0,174O0,484P0,00816S0,00141
Biomass P. thermotolerans                    biom-Pt          CH1,63N0,159O0,478P0,00502S0,00138
Biomass P. ethanolica				biom-Pe		    C1H1,63N0,165O0,454P0,00724S0,00137
Biomass S. cerevisiae                        biom-Sc          CH1,63N0,163O0,467P0,00753S0,00133
Biomass S. dispora				biom-Sd		    C1H1,62N0,180O0,452P0,00911S0,00144
carbamoyl phoshate                           CARP               CH2NO5P-2
CARBOHYDRATES                                CARBHYD            CH1,67O0,833
Carbon dioxide                               CO2                CO2
CDP                                          CDP                C9H12N3O11P2-3
chorismate                                   CHO                C10H8O6-2
cis-aconitate                                ACO                C6H3O6-3
citrate                                      CIT                C6H5O7-3
citydine diphosphate-diacylglycerol          CMP-DGOH           C45,3H78,7N3O15P2-2
CMP                                          CMP                C9H12N3O8P-2
CoA                                          CoA                C21H32N7O16P3S-4
CTP                                          CTP                C9H12N3O14P3-4
cystein                                      CYS                C3H7NO2S
dihydrofolate                                DHF                C19H19N7O6-2
dihydroxyacetone-phosphate                   DHAP               C3H5O6P-2
DNA                                          DNA                CH1,26N0,378O0,612P0,102
E4P                                          E4P                C4H7O7P-2
ETOH                                         ETOH               C2H6O
FAT                                          FAT                C52,9H97,5O6
formyl-THF                                   FTHF               C20H21N7O7-2
fructose 1,6-bisphosphate                    F16P               C6H10O12P2-4
fructose 6-phosphate                         F6P                C6H11O9P-2
fumarate                                     FUM                C4H2O4-2
GDP                                          GDP                C10H12N5O11P2-3
glucose 6-phosphate                          G6P                C6H11O9P-2
glutamate                                    GLM                C5H8NO4-1
glutamine                                    GLN                C5H10N2O3
glyceraldehyde 3-phosphate                   GAP                C3H5O6P-2
glycerol 3-phosphate                         GOH3P              C3H7O6P-2
glycine                                      GLY                C2H5NO2
glyoxylate                                   GLYO               C2HO3-1
GMP                                          GMP                C10H12N5O8P-2
GTP                                          GTP                C10H12N5O14P3-4
histidine                                    HIS                C6H10N3O2+1
homocitrate                                  HCIT               C7H7O7-3
homocysteine                                 HCYS               C4H8NO2S-1
homoserine                                   HSER               C4H9NO3
hydrogen                                     H                  H+1
IMP                                          IMP                C10H11N4O8P-2
isocitrate                                   ICIT               C6H5O7-3
isoleucine                                   ILE                C6H13NO2
isopropylmalate                              IPM                C7H10O5-2
leucine                                      LEU                C6H13NO2
LIPID                                        LIPID              CH1,87N0,00953O0,149P0,00953
lysine                                       LYS                C6H15N2O2+1
malate                                       MAL                C4H4O5-2
malonyl-CoA                                  MACoA              C24H33N7O19P3S-5
metal                                        metal              X
metheonine                                   MET                C5H11NO2S
methylene-THF                                METHF              C20H21N7O6-2
methyl-THF                                   MYTHF              C20H23N7O6-2
NAD                                          NAD                +1
NADH                                         NADH               H
NADP                                         NADP               +1
NADPH                                        NADPH              H
NH4                                          NH4                H4N+1
oleoyl-CoA                                   OLE-CoA            C39H64N7O17P3S-4
ornithine                                    ORN                C5H13N2O2+1
oxaloacetate                                 OXACT              C4H2O5-2
oxoadipate                                   OAD                C6H6O5-2
oxobutyrate                                  OBU                C4H6O3
oxoglutarate                                 OGL                C5H4O5-2
oxoisovalerate                               OIV                C5H7O3-1
oxygen                                       O2                 O2
palmitic acid                                PLM                C16H31O2-1
palmitoleoyl-CoA                             PLLM-CoA           C37H60N7O17P3S-4
palmityl-CoA                                 PLM-CoA            C37H62N7O17P3S-4
phenylalanine                                PHE                C9H11NO2
phosphate                                    Pi                 HO4P-2
phosphatidate                                PHD                C36,3H66,7O8P-2
phosphatidyl-choline                         PHD-CHO            C41,3H79,7NO8P
phosphatidyl-ethanolamine                    PHD-ETA            C38,3H73,7NO8P
phosphatidyl-serine                          PHD-SER            C39,3H73,7NO10P
phosphoenol-pyruvate                         PEP                C3H2O6P-3
phosphoribosyl-formamido-imidazole-carboamide    AICAR          C9H13N4O8P-2
proline                                      PRO                C5H9NO2
PROTEIN                                      PROT               CH1,58N0,275O0,318S0,00265-0,00587
PRPP                                         PRPP               C5H8O14P3-5
pyrophosphate                                PPi                O7P2-4
pyruvate                                     PYR                C3H3O3-1
quinone                                      Q     	           C
quinol                                       QH2                CH2
RIB5P                                        RIB5P              C5H9O8P-2
RIBU5P                                       RIBU5P             C5H9O8P-2
RNA                                          RNA                CH1,23N0,389O0,737P0,105
s-adenosyl-homocysteine                      SAH                C14H20N6O5S
s-adenosylmetheonine                         SAM                C15H23N6O5S+1
SED7P                                        SED7P              C7H13O10P-2
serine                                       SER                C3H7NO3
stearate                                     STE                C18H35O2-1
stearoyl-CoA                                 STE-CoA            C39H66N7O17P3S-4
succinate                                    SUC                C4H4O4-2
succinate-COA                                SUCCOA             C25H35N7O19P3S-5
sulfide                                      H2S                H2S
sulphate                                     SO4                O4S-2
tetrahydrofolate                             THF                C19H21N7O6-2
threonine                                    THR                C4H9NO3
tryptophane                                  TRP                C11H12N2O2
tyrosine                                     TYR                C9H11NO3
UDP                                          UDP                C9H11N2O12P2-3
UMP                                          UMP                C9H11N2O9P-2
UTP                                          UTP                C9H11N2O15P3-4
valine                                       VAL                C5H11NO2
water                                        H2O                H2O
XYL5P                                        XYL5P              C5H9O8P-2
```

**Literature references**

Daran-Lapujade, P., Jansen, M. L. A., Daran, J. M., Van Gulik, W., De Winde, J. H., & Pronk, J. T. (2004). Role of Transcriptional Regulation in Controlling Fluxes in Central Carbon Metabolism of Saccharomyces cerevisiae: A chemostat culture study. Journal of Biological Chemistry, 279(10), 9125–9138. <https://doi.org/10.1074/jbc.M309578200>

Roels, J. A. (1980). Simple model for the energetics of growth on substrates with different degrees of reduction. Biotechnology and Bioengineering, 22(1), 33–53. <https://doi.org/10.1002/bit.260220104>

Stock, D., Leslie, A. G. W., & Walker, J. E. (1999). Molecular architecture of the rotary motor in ATP synthase. *Science*, *286*(5445), 1700–1705. <https://doi.org/10.1126/science.286.5445.1700>

Verduyn, C., Stouthamer, A. H., Scheffers, W. A., & Van Dijken, J. P. (1991). A theoretical evaluation of growth yields of yeasts. *Antonie van Leeuwenhoek*, *59*, 49–63.
